# Dynamic causal communication channels between neocortical areas

**DOI:** 10.1101/2021.06.28.449892

**Authors:** Mitra Javadzadeh, Sonja B. Hofer

## Abstract

Dynamic pathways of information flow between distributed brain regions underlie the diversity of behaviour. However, it remains unclear how neuronal activity in one area causally influences ongoing population activity in another, and how such interactions change over time. Here we introduce a causal approach to quantify cortical interactions by pairing simultaneous electrophysiological recordings with neural perturbations. We found that the influence visual cortical areas had on each other was surprisingly variable over time. Both feedforward and feedback pathways reliably affected different subpopulations of target neurons at different moments during processing of a visual stimulus, resulting in dynamically rotating communication dimensions between the two cortical areas. The influence of feedback on primary visual cortex (V1) became even more dynamic when visual stimuli were associated with a reward, impacting different subsets of V1 neurons within tens of milliseconds. This, in turn, controlled the geometry of V1 population activity in a behaviourally relevant manner. Thus, distributed neural populations interact through dynamically reorganizing and context-dependent communication channels to evaluate sensory information.

## Introduction

Animal behaviour arises from distributed patterns of neural activity across many brain regions. Neural activity in any one region does not develop in isolation, but is constantly shaped by, and in turn shaping, activity in others. Hence, brain-wide activity patterns underlying behaviour evolve on a moment-to-moment basis through the continuous exchange of information between brain areas. Understanding such complex networks of information flow requires looking beyond the anatomical connections that provide the substrate for communication, and asking how distributed brain areas dynamically influence each other^1–3^.

Statistical relations of activity across brain regions have been traditionally used to infer causal interactions between them^4–12^. The majority of previous studies have relied on macroscopic measures of neural activity, such as the local field potential, EEG or fMRI BOLD signals^13–19^. Such one-dimensional summary signals reflect combined activity from many neurons. However, several recent studies have emphasized that specific patterns of population activity, rather than the average level of activity in an area, are crucial for how it relates to the activity of downstream targets^20–22^. For instance, in the motor cortex of macaques only certain activity patterns are predictive of muscle movement, while other patterns fall into a ‘null space’ and are not related to movement^20^. Likewise, in the macaque visual cortex, only a small subset of population activity patterns are statistically related to downstream activity^21^. Such findings have highlighted the importance of single cell resolution measurements in characterizing interareal interactions and have spurred the development of diverse multivariate statistical analysis methods to quantify the interactions between populations of neurons recorded across different brain areas^4–6^.

However, despite the recent advances in multi-area population recording and analysis methods, the main challenge in quantifying inter-areal communication remains unresolved: correlated patterns of activity measured across two brain areas could arise from causal influences these areas have on each other, or alternatively, could be due to common or correlated inputs to both areas, and therefore not reflective of causal interactions. It has been shown that statistical approaches cannot distinguish these possibilities^4,23–25^. This problem is particularly pronounced in highly interconnected networks with a large number of unobserved sources of variability, such as other brain regions which were not recorded^24^. Therefore, while previous studies have shown that the statistical relations of simultaneously recorded neural activity in neocortical areas are modified depending on behavioural demands^1,15–18,26,27^, we do not yet know if this is reflective of changes in causal influences between areas.

Overcoming this long-acknowledged obstacle in the field requires a causal approach to measuring inter-areal interactions, based on manipulations of neural activity. Optogenetic manipulations have previously been used to examine the effect of silencing or activating different brain areas on long-range targets^28–30^. However, such studies have not focused on how communication between populations of neurons is dynamically organized, or how it causally shapes neural activity patterns at different time points or in different behavioural contexts.

Accordingly, the principles of how distributed neuronal populations communicate are still unclear. Specifically, it remains unresolved how the patterns of influence from one area on its target populations are structured, whether these causal influences are static or change over time, and whether communication channels between areas can be dynamically regulated depending on the behavioural state of the animal.

## Results

### Measuring causal interactions between neocortical areas

To address these questions, we developed a paradigm for measuring causal interactions between cortical areas by manipulating neuronal activity and characterizing inter-areal communication on the neuronal population level with single-cell resolution. Using direct experimental manipulations to establish causal interactions eliminated the need for relying on statistical associations of neural activity. We measured the directional influence of one cortical area (source) on another (target), focusing on feedforward and feedback influences between primary visual cortex (V1) and lateromedial higher visual area (LM) in mice (**Fig. 1a,b**). We performed simultaneous electrophysiological multi-channel recordings at retinotopically matched locations in the two areas (**Extended Data Fig. 1a-c**). We then silenced one of the two areas for brief time windows using optogenetic activation of inhibitory parvalbumin-expressing interneurons expressing channelrhodopsin-2 (ChR2)^31,32^ and measured the instantaneous effect on the recorded neurons in the target area, revealing the causal influence of the source area on the target neurons’ activity.

**Fig. 1.**
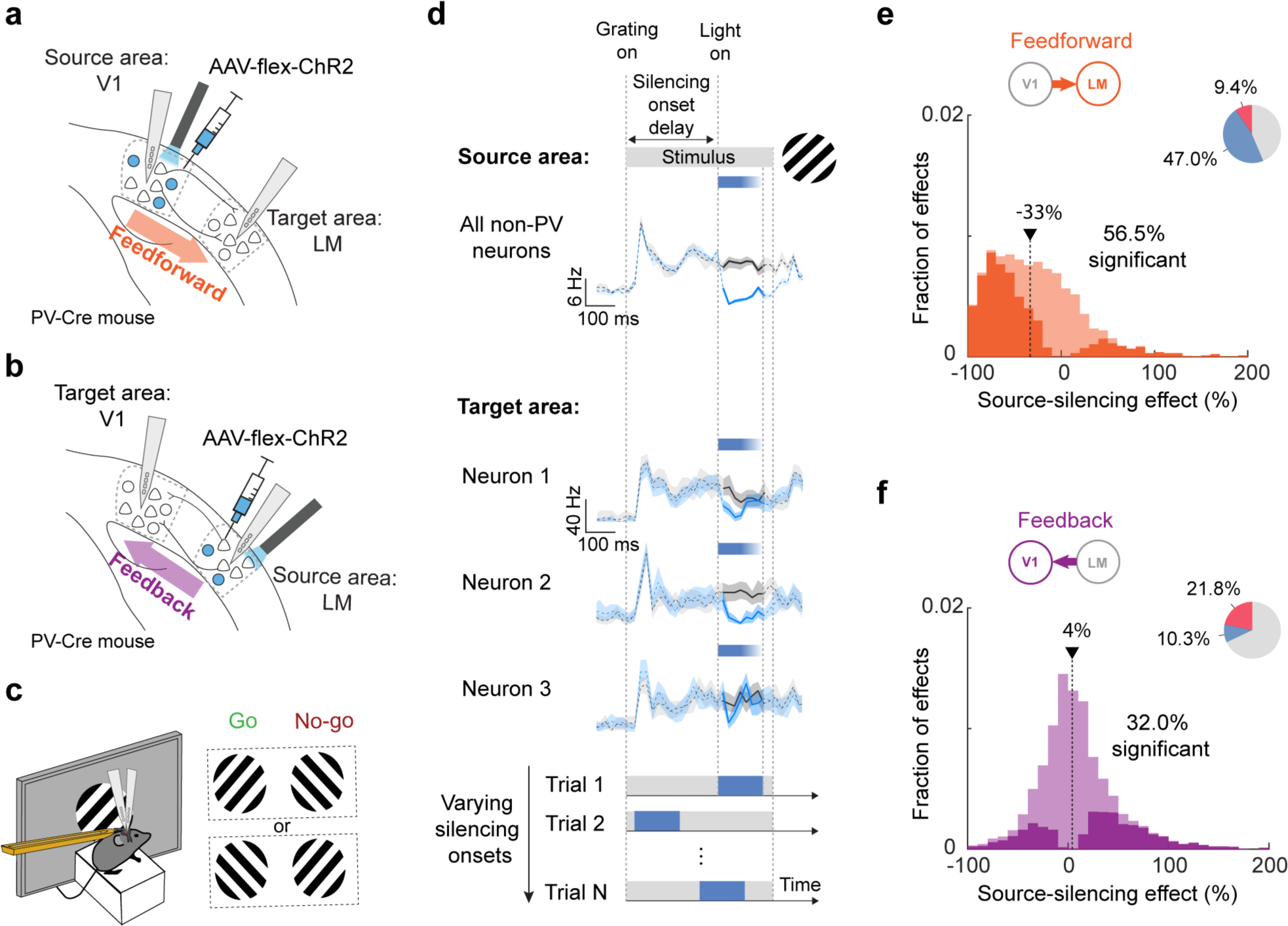
Experimental design for measuring causal influences between V1 and LM. **a,** Paired recordings in retinotopically matched regions of V1 and LM with silicon probes in PV-Cre mice. In order to measure the influence of V1 on LM cells (feedforward influence), V1 was silenced through light-mediated activation of parvalbumin-expressing inhibitory cells expressing ChR2 after injection of AAV-flex-ChR2. **b,** as in **a,** but with optogenetic silencing of LM in order to assess influence of LM on V1. **c,** Recordings and optogenetic manipulation were performed during a Go/No-go task. Head-fixed, stationary mice were presented with two differently orientated stationary grating stimuli (45 or -45 degrees), only one of which was rewarded. The identity of the rewarded stimulus was randomized across mice. Mice reported the rewarded stimulus by licking a spout, which triggered the delivery of the reward. **d,** Example traces of spiking activity during V1 silencing. The traces denote stimulus-evoked activity in the source (V1) and target (LM) area in control (black traces) and silencing trials (blue traces), binned at 20 ms. Shading is the 95% bootstrapped confidence interval. The grey bar on top indicates the presence of the visual grating stimulus (45 or -45 degree orientation, 40 degrees diameter). Blue bars indicate the time of optogenetic silencing. Top, average activity of all non-PV neurons (broad action potential waveform) in the source area (V1). Middle, average activity in control and silencing trials of three example neurons in the target area (LM). Bottom, dynamic characterization of inter-areal influence by varying silencing onset times. For each trial, the silencing onset time is chosen, in a randomized manner, from a pool of 8 values tiling the stimulus duration with ∼65ms resolution. **e,** Distribution of the effects of silencing V1 on the firing rate of LM neurons from all silencing time windows and stimuli (feedforward influences, n = 7 mice, n = 278 neurons, 8 silencing time windows), calculated as percentage change in firing rate when silencing V1. Negative source-silencing effects indicate decreases in firing rate during silencing, and therefore a net excitatory influence of V1 on the LM cell. Positive source-silencing effects indicate increases in firing rate, and therefore a net inhibitory influence of V1 on the LM cell. Significant effects are shown in bright colours (56.5 % of all effects). The arrow denotes the median of the distribution. The pie chart shows the fraction of significant increases (red) and decreases (blue) in LM neuron firing rates during V1 silencing. **f,** As in **e,** but distribution of the effects of silencing LM on the firing rate of V1 neurons (feedback influences, n= 7 mice, n = 242 neurons, 8 silencing time windows, 32.0 % of all effects significant).

In order to assess how behavioural state modulates cortical communication, we trained mice to perform a Go/No-go task. Animals learned to discriminate two stationary visual grating stimuli of which only one was rewarded (**Fig. 1c**, **Extended Data Fig. 1d,e**). Optogenetic silencing was performed in 150 ms time windows during the presentation of the 500 ms long grating stimuli. The onset of silencing varied in each trial, tiling the duration of the visual stimulus in ∼65ms steps (**Fig. 1d**). The effect of silencing on target area firing rates was only assessed during the time window of optogenetic manipulation. This approach allowed us to quantify how the influence of the source on the target area may change over time during cortical processing of behaviourally-relevant stimuli with a resolution of tens of milliseconds (**Fig. 1d**).

### Inter-areal influences are diverse and vary with time

We first characterized the average influence of V1 activity on LM neurons (feedforward influence), irrespective of silencing time window and task condition. We measured the trial-averaged percentage of change in the firing rate of LM neurons during each time window in which V1 was silenced. A large fraction of LM neurons was suppressed by V1 silencing (**Fig. 1e**, median change in spiking rate = -33%, with 54.9 ± 7.6 %, mean ± s.e.m of neurons significantly affected in at least one time window), confirming a predominantly excitatory feedforward influence of V1 on this higher visual area. The extent to which LM neurons were suppressed when V1 was silenced was partly related to their average firing rate, cortical depth at which they were located and their relative receptive field position, according to a multivariate regression model (**Extended Data Fig. 2**, see Methods). Specifically, LM neurons were less affected by V1 activity when they resided in deep cortical layers^33^ or when their spatial receptive field was far displaced from the retinotopic location of the centre of optogenetic manipulation in V1 (**Extended Data Fig. 2d,e**), revealing a structured influence of feedforward input.

**Fig. 2.**
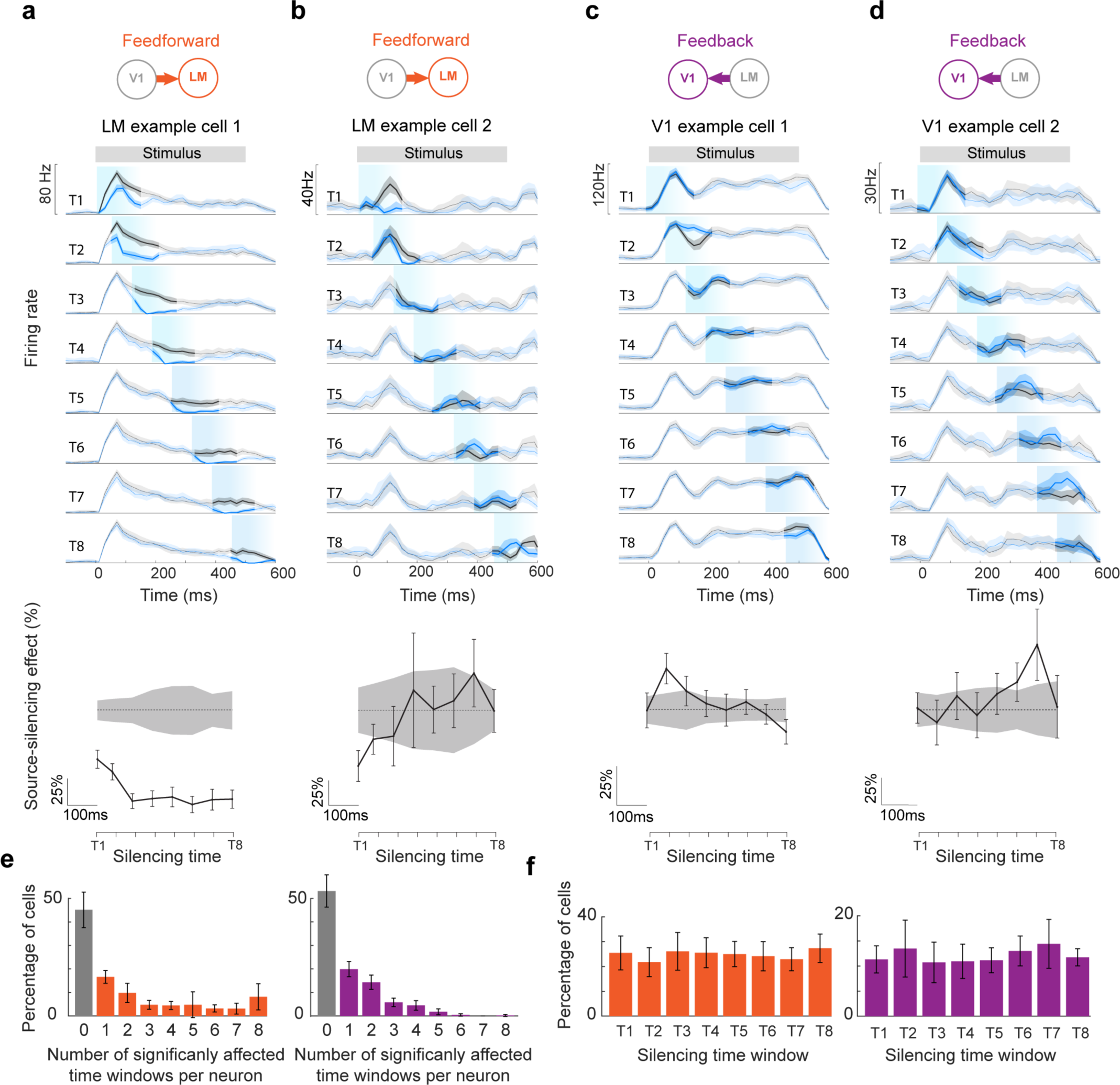
Temporal profile of feedforward and feedback influences on example target neurons in V1 and LM. **a,b,** Top, firing rates in control (black), and V1 silencing (blue) trials during Go stimuli at 8 different silencing onset times (T1-T8). The grey bar on top indicates the presence of the visual grating stimulus. Light blue shading marks the timing and duration of optogenetic silencing. PSTHs were smoothed over 60 ms using a moving average filter. Shading indicates the 95% bootstrapped confidence intervals of the mean. Bottom, feedforward source-silencing effects on the target neuron over time, calculated as percentage change in firing rate when silencing the source area. Zero is indicated by a dashed line. Chance levels are indicated by grey shading, calculated as 95% confidence interval of percentage differences in the firing rate of bootstrapped control trials. Error bars indicate 95% bootstrapped confidence interval of the mean. **c,d,** As in **a,b,** but showing effects of silencing LM on two example V1 cells. **e,** Distribution of the number of time windows in which individual neurons were significantly affected (after Bonferroni correction for 8 comparisons) by V1 silencing (left, LM neurons, n = 7 mice, 278 neurons) and by LM silencing (right, V1 neurons, n = 7 mice, 242 neurons). Data was averaged across Go and No-go trials. Error bars depict 95% confidence interval (2 x s.e.m) of the mean across animals. **f,** Percentage of neurons showing a significant change of firing rate during each silencing time window for LM neurons during V1 silencing (left, one-way ANOVA, p = 0.9) and for V1 neurons during LM silencing (right, one-way ANOVA, p = 0.8). Error bars as in **e**.

We then performed equivalent experiments while silencing higher visual area LM to measure the effect of LM feedback on the activity of V1 neurons. Silencing LM changed the firing of a substantial number of V1 neurons (**Fig. 1f**; 46.9 ± 6.9 % of neurons significantly affected in at least one time window), but did not have a clear net excitatory or inhibitory influence (median change in spiking rate of 4%, **Fig. 1f**). Instead, LM activity had diverse effects on individual V1 neurons, including decreased or increased activity upon LM silencing. A previous study in awake primates found similarly diverse excitatory and inhibitory influences of higher visual area V2 on V1^34^. In contrast to the feedforward effect of V1 on LM, we did not identify any attributes of V1 neurons that could explain by how much or in which direction they were affected by feedback from LM (**Extended Data Fig. 2c,f,g**). The average effect of silencing feedforward and feedback inputs was similar during the rewarded grating stimulus (Go trials) and the non-rewarded stimulus (No-go trials, **Extended Data Fig. 3**, Go vs No-go feedforward influence p = 0.87, feedback influence p = 0.78, two-sided Wilcoxon rank-sum test).

**Fig. 3.**
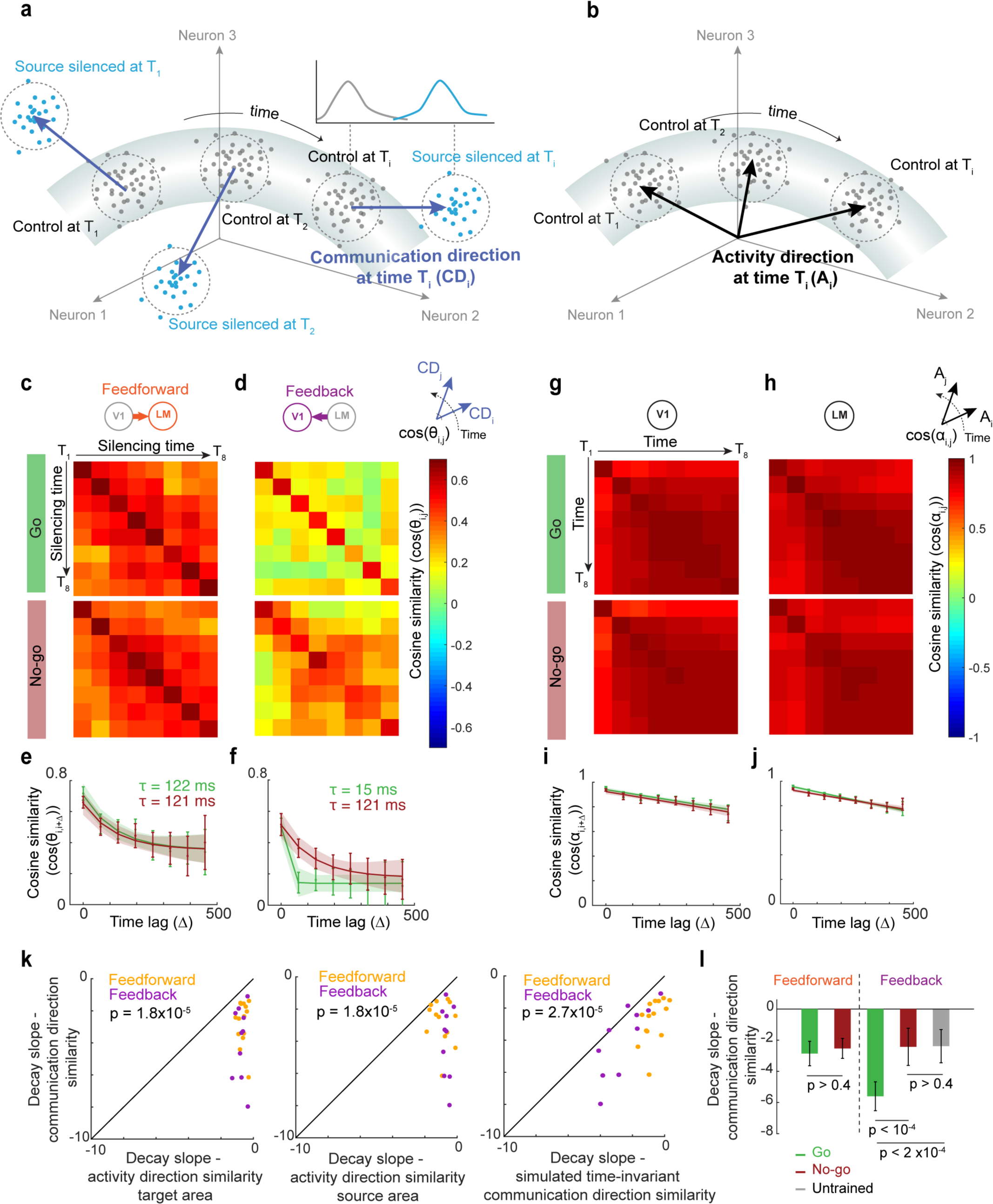
Population-level dynamics of inter-areal influences. **a,** Schematic of the population analysis framework. Each point in the neural activity space specifies the population activity in the target area in a given time window (150 ms) of one trial. Grey points denote control trials and blue points source-silencing trials. The communication direction (blue arrows) is calculated for each time window as the direction that maximally separated the activity in control trials (grey cloud of points) from that in source silencing trials (blue cloud of points, see methods). **b,** Activity directions (black arrows) are calculated for each time window as the direction specifying the centre of the trial-averaged activity in control trials in that time window (centre of the grey cloud of points). **c,** Matrices depicting cross-validated cosine similarity between communication directions (as shown in **a**) of all pairs of silencing time windows during the Go (top) and No-go stimulus (bottom) for feedforward influences on LM (silencing of V1). Matrices are averaged across animals (N = 7 mice). **d,** As in **c**, but for the feedback influences on V1 (silencing of LM) (N = 6 mice). **e,** Cosine similarity of pairs of feedforward communication directions in different time windows as a function of the time lag between them during the Go (green) and No-go stimulus (red). Error bars depict the 95% confidence interval of the mean (2×s.e.m). Lines depict exponential fits, shading shows 95% prediction bounds of the fits. The decay time constants of the exponential fits (τ) are shown for Go (green) and No-go (red) stimuli. **f,** as in **e,** but for feedback communication directions. **g,** Matrices depicting cross-validated cosine similarity between activity directions (as shown in **b**) in V1 (n = 13 mice) of all pairs of time windows in control trials during the Go (top) and No-go stimulus (bottom). **h,** as in **g,** but showing the cross-validated cosine similarity of activity directions in LM (n = 13 mice). **i,** Pairwise cosine similarity of V1 activity directions in different time windows as a function of the time lag between them during the Go (green) and No-go stimulus (red). Error bars depict the 95% confidence interval of the mean (2×s.e.m). Lines depict linear fits, shading shows 95% prediction bounds of the fits. **j,** same as **i,** but for LM activity directions. **k,** Relationship between the initial slopes (slopes between lag 0 and lag 1) of the decay over time lags of communication direction similarities and activity direction similarities in the target area (left), and in the source area (middle), and between initial decay slopes of communication direction similarities in the real dataset and the simulated dataset assuming time-invariant inter-areal influences (right). P-values from two-sided Wilcoxon signed-rank tests. Orange and purple dots show data from individual animals during V1 silencing (feedforward influences) and LM silencing (feedback influences) experiments, respectively. **l,** Initial slopes (slopes between lag 0 and lag 1) of the decay over time lags of feedforward and feedback communication directions in Go (green) and No-go (red) trials of trained animals, and of feedback communication directions in untrained animals (grey). Error bars depict the 95% confidence interval of the mean (2×s.e.m). P-values from Wilcoxon two-sided signed-rank test for comparisons between Go and No-go in trained animals, and from two-sided Wilcoxon rank-sum test for comparisons with untrained animals.

Quantification of average source-silencing effects on the target area does not reveal how neurons are influenced at different moments in time. To determine how the two cortical areas influenced each other over the course of visual stimulus presentation, we compared how similarly target area activity was affected in eight different silencing time windows (T1-T8) tiling the visual stimulus duration (**Fig. 1d**, **Fig. 2**). In some target area neurons, the change in firing rate induced by source-area silencing was relatively similar during all time windows (**Fig. 2a**), revealing a constant influence of the source area on these neurons over time. By contrast, many neurons showed varying changes in their firing rates during different silencing windows (**Fig. 2b-e**). For instance, feedback from LM often exerted a significant effect on a V1 neuron only during a short time period (as short as < 100ms, **Fig. 2c-e**). These silencing effects were consistent from trial to trial for the same neuron, but not consistent between neurons, with individual cells affected at different times during the stimulus presentation (**Fig. 2f**).

### Fast re-structuring of inter-areal influences on population activity patterns

To quantify the diverse and temporally variable contributions of a source area to target neurons’ activity, we derived a population-level measure of inter-areal influence between V1 and LM. In a given time window, the population activity of *n* simultaneously recorded neurons in the target area can be described as a point in an *n*-dimensional space in which the coordinates specify the firing rate of each neuron. In this framework, one set of points represents the population activity during a specific time window in all control trials without optogenetic manipulation, and another set of points constitutes the population activity in all silencing trials in which source area activity was absent during the same time window (**Fig. 3a,b**). To find a relation between these population activity states, we identified the direction in the *n*-dimensional activity space along which the population activity during source-silencing trials could be maximally discriminated from control trials using linear discriminant analysis^35^ (see Methods, **Extended Data Fig. 4a-f**). This direction denotes a mode of target area activity (pattern of activity in a specific neural ensemble) which is causally dependent on the activity in the source area, and therefore is termed the causal communication direction (**Fig. 3a**).

**Fig. 4.**
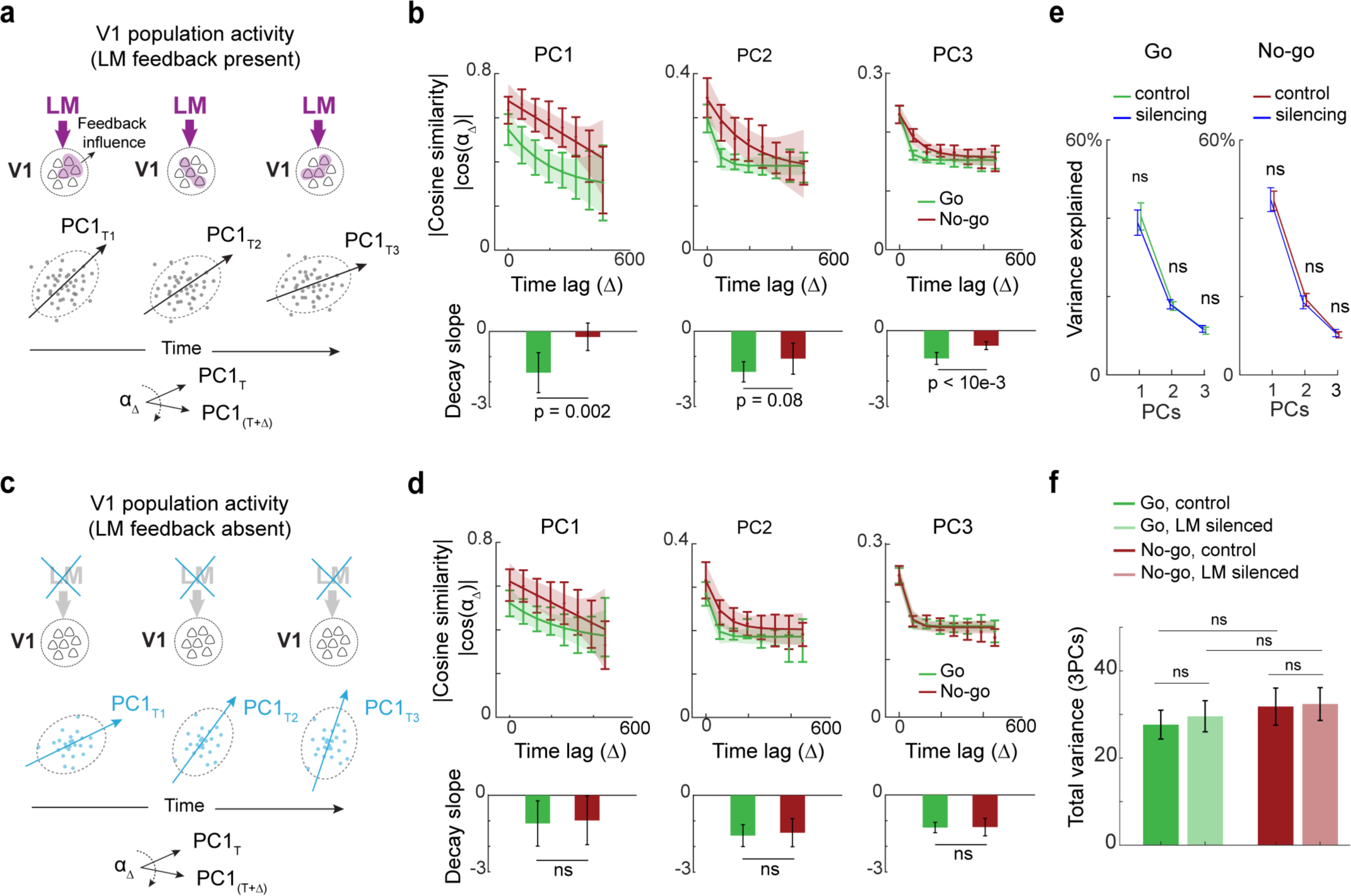
Feedback modulates the covariance structure of V1 population activity. **a,** Schematic of V1 population activity in different time windows in control trials. The cloud of points indicates the trial to trial variability in the patterns of V1 population activity. The shape of this cloud is described by the principal components. Principal components (PC directions) were calculated separately for each time window, and their similarity was compared over time. **b,** Top, Absolute value of pairwise cross-validated cosine similarity of the first (left), second (middle), and third (right) principal components of V1 activity in different time windows as a function of the time lag between them, in Go (green) and No-go (red) trials. Error bars depict the 95% confidence interval of the mean (2×s.e.m). Lines depict exponential fits, shading shows 95% prediction bounds of the fits. Bottom, initial slope (slope between lag 0 and 1) of the decay of principal component similarity over time lags for Go and No-go trials. Error bars depict the 95% confidence interval of the mean (2×s.e.m). P-values from two-sided Wilcoxon signed-rank tests. **c,** As in **a,** but during silencing of LM, depicting principal components of V1 population activity over time in the absence of LM feedback. **d,** As in **b,** but during LM silencing trials (in the absence of LM feedback to V1) **e,** Percentage of variance explained by each of the first three principal components in V1 during Go (green, left) and No-go (red, right) stimuli in control and silencing trials (blue) across all time windows. Error bars depict the 95% confidence interval of the mean (2×s.e.m). **f,** The sum of variance across the first three principal components in V1 for control and LM silencing trials, during the Go (green) and No-go (red) stimulus averaged across all time windows. Error bars depict the 95% confidence interval of the mean (2×s.e.m).

We determined causal communication directions separately for the eight time windows during the visual stimulus and quantified their pairwise similarity by calculating the cosine similarity between communication directions (the dot product of unit-length vectors) for each pair of silencing time windows. Cosine similarities were calculated in a cross-validated manner (see Methods), and therefore provided a robust estimate of similarity with respect to trial-to-trial variabilities. We visualized cross-validated cosine similarities between all possible pairs of communication directions of all silencing time windows T1 to T8 in cosine similarity matrices (**Fig. 3c,d**), and quantified how fast communication directions changed by comparing the similarity of directions over all time lags (**Fig. 3c-f**). This analysis allowed us to address how stable the causal communication direction was over time and thereby assess the time-varying causal influence neuronal populations in the two cortical areas had on each other.

The similarity of both feedforward and feedback communication directions decayed relatively rapidly with time (**Fig. 3e,f**, exponential decay time constant (τ) < 122 ms for all conditions), indicating that both feedforward and feedback influences on population activity in V1 and LM are temporally dynamic. To test if this time-varying influence can simply be explained by temporally variable population activity in the source or target area, we compared how the patterns of population activity within V1 and LM changed over time. The trial averaged population activity at each time window was quantified using population activity directions, defined as the centre coordinate of the set of points in the population activity space corresponding to activity in control trials at that time window (**Fig. 3b**). We calculated cross-validated cosine similarities between the activity directions of control trials of all eight time windows (**Fig. 3g-j**). The modes of activity in both V1 and LM were very similar across different time windows, and the similarity of activity directions therefore decayed much more slowly over time than the similarity of both the feedforward and feedback communication directions (**Fig. 3k**, left and middle, all p-values < 2 × 10^-5^, two-sided Wilcoxon signed-rank test). This was the case even if only the 20% of source area neurons with the most time-varying activity were included for calculating activity directions (**Extended Data Fig. 4g-n**, see Methods). Therefore, the fast changes in communication directions cannot be explained by a subset of source area neurons with particularly high temporal variations in their firing rate providing input to the target area.

However, the above analyses rely on comparisons between LDA vectors capturing manipulation effects and population activity vectors, which are different measures, with different levels of robustness to trial-to-trial noise (cross-validated similarity at time lag 0, see Methods). To corroborate that these differences cannot explain the faster decay in the similarity of communication directions compared to activity directions, we generated an artificial dataset to simulate how similar communication directions of different time windows would be if source-silencing effects on individual neurons were invariant in time. For each target neuron, we used its experimentally measured activity in control trials and simulated the activity in source-silencing trials, such that the effect of silencing was the same in each time window while maintaining the experimentally observed distribution of silencing effects per cell (see Methods). This allowed us to directly compare changes in communication directions with and without time-varying influences of silencing, while preserving measured neuronal activity dynamics and the diversity of single-cell silencing effects, as well as experimentally observed trial-to-trial variability of firing rates. The communication directions of this simulated dataset changed slower over time than those of the experimental data (**Fig. 3k**, right, p = 2.7 x 10^-5^, two-sided Wilcoxon signed-rank test, **Extended Data Fig. 5a,b**), confirming that the fast decay in the similarity of communication directions cannot be explained by trial-to-trial noise or the temporal dynamics of neuronal activity patterns. Therefore, our data show that communication between cortical areas is dynamic and reorganizes over time, such that different ensembles of target area neurons are engaged by the source area at different time points during processing of a visual stimulus.

### Temporal dynamics of feedback influences depend on behavioural relevance

Previous studies have shown that the relationship between neuronal activity in different cortical areas during processing of a sensory stimulus is modulated by the relevance and behavioural consequences of the stimulus^17,18,26,36,37^. These results raise the possibility that information flow between areas can be flexibly adjusted by the behavioural context of sensory signals. To determine if the causal influence of areas V1 and LM on each other changes depending on behavioural task demands, we compared how the time-varying effects of feedforward and feedback input on population activity differed in Go and No-go trials during the visual discrimination task. Only the Go stimulus was associated with a reward and therefore behaviourally most relevant.

The temporal dynamics of feedforward communication directions, denoting how much the activity mode in LM that was causally dependent on activity in V1 changed over time, were similar during the Go and No-go stimulus (**Fig. 3e,l**, exponential decay time constant, Go trials: 122 ms, 81 to 153 ms 95% confidence interval, CI; No-go trials: 121 ms, 89 to 170 ms CI; p = 0.99, permutation test). In contrast, the dynamics of feedback influences were much faster during the Go than the No-go stimulus, with the cosine similarity of feedback communication directions from LM to V1 decaying at a time constant of ∼ 15 ms only during the Go stimulus (**Fig. 3f,l**, exponential decay time constant, Go trials: 15 ms, 0 to 46 ms CI; No-go trials: 121 ms, 93 to 163 ms CI; p = 1.3 × 10^-4^, permutation test). These effects could not be explained by differences in V1 or LM activity, since the dynamics of population activity directions in both areas were similar during Go and No-go trials (**Fig. 3g-j**, **Extended Data Fig. 5c**), as was the level of trial-to-trial variability of communication directions (**Fig. 3f** time lag 0; see Methods; Go vs No-go trials p > 0.5 two-sided Wilcoxon signed-rank test). To rule out that differences in feedback communication could be due to differences in motor actions during Go and No-go trials such as licking, we compared the dynamics of feedback influences during an early epoch before mice responded to the visual stimulus (50-200ms post stimulus onset) and a late epoch in which mice showed lick responses to the Go stimulus (> 250ms post stimulus onset). In both epochs, changes in feedback communication directions were faster during Go stimulus trials (**Extended Data Fig. 5d**).

These data indicate that feedback communication during sensory processing is regulated by behavioural relevance. Accordingly, we found that in untrained mice that were passively viewing the visual stimuli, feedback communication directions exhibited temporal dynamics similar to those during No-go trials in trained animals (**Fig. 3l**, **Extended Data Fig. 5e**). Hence, the ensembles of V1 neurons influenced by LM activity changed much more rapidly when the visual stimulus processed by these cortical areas is predictive of reward (see also **Extended Data Fig. 5f**). While we did not find similar effects of stimulus relevance on the temporal patterns of feedforward communication, the influence of V1 onto downstream targets may be modulated by other behavioural variables not captured by our task, such as spatial attention ^17,18,26^.

### Feedback restructures V1 covariance patterns depending on behavioural relevance

How does the dynamic influence of feedback impact stimulus coding in V1? Since trial-averaged V1 activity directions changed at similar rates during the stimulus in Go and No-go trials (**Fig. 3i,j**, **Extended Data Fig. 5c**), we looked beyond average activity patterns and examined the geometry of trial-to-trial fluctuations. Neurons in cortical areas form functional subnetworks within which activity is correlated from trial to trial. The organization of these co-fluctuations - the covariance structure of population activity - has important implications for the coding capacity of and the readout of information from an area^38–40^. These co-fluctuations can be captured by principal component analysis, wherein the first few principal components (PCs) define the most prominent modes of correlated activity in V1. We computed the first three PCs of V1 population activity (explained variance = 72 ± 9.2%, mean ± s.d.) separately for the eight time windows during visual stimulus presentation, and used cross-validated cosine similarity to compare their differences over time as described above (see Methods).

Interestingly, covariance patterns of V1 activity were not stable but appeared to re-organize over time, as the dominant PCs changed their direction in succeeding time windows (**Fig. 4a,b**). Moreover, the PCs changed direction at a faster rate in Go compared to No-go trials, revealing that functional V1 subnetworks re-organized faster during presentation of the behaviourally relevant Go stimulus. However, this was only the case in control trials when LM feedback was intact (**Fig. 4a,b**). During silencing trials, when feedback from LM was absent, the PCs’ temporal evolution was similar in Go and No-go trials (**Fig. 4c,d**), indicating that the temporal evolution of V1 covariance patterns depended on feedback from higher visual area LM. In contrast, how much of the variance of V1 activity each PC could explain and their total variance was not different in Go and No-go trials and was not changed when LM was silenced (**Fig. 4e,f**, see Methods), indicating that feedback only rotated the covariance structure, without affecting the overall amount of trial-to-trial variability in V1 population activity.

Given that feedback regulates the correlation structure of V1 population activity, we asked how these influences were reflected in the temporal and pairwise dependencies of firing rates at the single neuron level. Temporal dependencies in the firing rate of individual V1 neurons over time, quantified by the autocorrelation function over different time lags, was also different during Go and No-go stimuli, as autocorrelations were lower during Go stimuli (**Extended Data Fig. 6a**). This difference was absent when feedback from LM was silenced (**Extended Data Fig. 6a**). Moreover, V1 neurons whose firing rate was more strongly affected by LM silencing showed faster decay of autocorrelations (**Extended Data Fig. 6b**), confirming that the decreased dependency of neuronal firing rates with time was related to feedback from LM. Finally, pairwise noise correlations between V1 neurons were on average weaker in Go than in No-Go trials (**Extended Data Fig. 6c**), confirming previous findings of decreased firing rate interdependencies in cortical areas during processing of behaviourally relevant visual stimuli^41^. However, feedback from LM did not affect the average strength of pair-wise noise correlations nor their modulation by task demands (**Extended Data Fig. 6c,d**).

Therefore, feedback from LM affects specific aspects of V1 population activity patterns. Feedback renders correlation patterns in neuronal firing rates more dynamic over time without affecting the overall variability of population activity, and therefore contributes to the continual restructuring of functional V1 subnetworks during stimulus coding. Moreover, the rate at which V1 functional subnetworks reorganize is modulated by behavioural demands through a change in the temporal dynamics of feedback influences from LM.

## Discussion

By using a causal approach to study inter-areal communication in the neocortex, we quantified the dynamics of feedforward and feedback interactions between cortical visual areas. We found that the influence of one area on another is specific - only subsets of target neurons are affected at each moment in time, and highly dynamic - different subsets of neurons can be affected at different moments in time within tens of milliseconds. This was in stark contrast to the stable activity patterns elicited by visual stimulation within the areas themselves. Moreover, these effects are modulated by the animal’s behavioural state.

Previously, specific patterns of population activity have been shown to be important for communication between areas. Activity in one area can be rendered effective or ineffective in influencing a target area depending on the degree by which it matches specific activity patterns, thus forming inter-areal communication subspaces^20–22^. Our findings indicate that these subspaces dynamically re-organize over short time scales relevant for stimulus processing and are subject to modulation by behavioural state.

Which mechanisms could underlie dynamic communication on the fast time scales observed in this study? Our method measures effective connectivity between areas^1^. The described effects could therefore be mediated by mono-synaptic intracortical projections as well as poly-synaptic pathways, for instance through the thalamus^42,43^. While our recordings did not provide selective access to target-projecting neurons in the source area, the population activity patterns of even the most time-varying source area neurons could not account for the fast dynamics of inter-areal influences (**Extended Data Fig. 4g-n**). Inter-areal communication could be regulated by changes in the synchrony of action potential timing within or between areas^11,12,44,45^. Neural activity oscillations have been proposed to enable such changes in synchrony^12,46,47^. However, it is unclear how the fast dynamics in feedback influences could be achieved by long time constants required for resonance or entrainment of oscillations^19,47^. Alternatively, subpopulations of target area neurons could be rendered more or less susceptible to inter-areal influences in specific moments in time by local circuit mechanisms, nonlinear amplification, or additional long-range input, for instance from higher-order thalamic nuclei such as the pulvinar^48^.

We found that causal feedback communication was modulated by the animals’ behavioural state, as the patterns of LM feedback influence on V1 varied much faster over time during the visual stimulus that was associated with a reward and therefore behaviourally most relevant. Cortical feedback projections have been suggested to influence sensory processing in various ways^34,49–56^, but their function is still unclear. We found that feedback from LM can alter dependencies in V1 single cell and population activity patterns over time. At the single cell level, we examined temporal and spatial dependencies in V1 using autocorrelations and pairwise noise correlations. Interestingly, feedback selectively modulated the time scales of autocorrelations in V1, without affecting the average strength of pairwise noise correlations. Previous studies have reported reduced noise correlations in attentive states, hypothesized to increase the coding capacity during attention, and thought to be mediated by top-down input^41,57^. We observed on average lower noise correlations during the rewarded Go stimulus, consistent with its higher behavioural relevance. However, our findings do not support a role of feedback from higher visual areas in this state-dependent modulation of noise correlations in V1, since LM feedback had no effect on noise correlation strength in either the Go or the No-go stimulus condition. Feedback may instead regulate computations in V1 by controlling the time scales of visual processing. Previous studies have shown a diversity of spike rate autocorrelation time constants across different brain areas^58,59^. This diversity potentially reflects area-specific temporal specializations for computations that require input integration on different time scales^58–60^. For instance, short time scales could be beneficial for rapid detection of stimuli, while longer time scales may support computations that depend on longer integration of signals. Our findings show that these time constants can be dynamically modulated by feedback to optimize V1 circuits according to the task demands.

At the population level, we found that feedback modified the geometry of the V1 covariance structure in a context-dependent manner^61^ by rotating the principal components of V1 activity without changing the variance along each principal component. The covariance structure of neural activity in a network describes the patterns of correlated variability between all recorded neurons and is therefore indicative of the functional subnetworks within an area. The structure of these subnetworks plays an important role in shaping the neural code and the readout of information from an area^38–40^. Since the covariance structure relates to the connectivity between neurons^38,62^, it is usually considered a static property of networks. However, we found the structure of correlated variability to be surprisingly dynamic: The most prominent modes of variability in the network (the principal components) rotated over time, leading to temporal restructuring of functional subnetworks in V1. This may enable V1 populations to conduct different computations at different moments in time during visual processing. Importantly, the speed of these changes in the covariance structure was modulated by cortical feedback in a context-dependent manner. Together, our findings therefore indicate a novel role of cortical feedback in temporally organizing the computations in V1.

Feedback projections are thought to be important for perceptual inference^63–65^. Several studies have suggested that populations of neurons can perform efficient Bayesian inference using Markov Chain Monte Carlo (MCMC) sampling algorithms, where the firing of each neuron represents stochastic samples from the posterior distribution (a probabilistic representation of the beliefs about sensory information given the input)^66,67^. MCMC algorithms produce autocorrelated samples which allow sampling from high dimensional distributions, but could lead to slow inference. Consequently, the efficiency of these samplers could be modulated by controlling the time constant of their autocorrelation function^67^. LM feedback selectively modulates temporal interdependencies (autocorrelations) in V1 circuits, and may thereby play a role in optimizing probabilistic sampling processes according to behavioural demands. Moreover, feedback from higher cortical areas has been proposed to provide expectations and beliefs about the world to earlier processing stages during perceptual inference^49,63–65^. The rapid changes in the influence of feedback on V1 activity over time may be a hallmark of evolving predictions in a dynamically changing environment.

In summary, we find that cortical areas interact through dynamically rotating communication channels to process and disambiguate sensory signals depending on the behavioural context.

## Methods

### Mice

All experiments were conducted in accordance with institutional animal welfare guidelines and licensed by the UK Home Office and the Swiss cantonal veterinary office. A total of 20 PV-Cre mice^68^ were used. Electrophysiological recordings and optogenetic manipulations were performed in 4 untrained mice and 16 mice trained in the visual discrimination task (7 and 9 mice with silencing of V1 and LM, respectively, 2 out of 16 mice were excluded due to their task performance, see analysis of behaviour). Mice were of either gender and were between 6 and 14 weeks old at the start of the experiment.

### Surgical procedures and virus injection

Prior to surgery, mice were injected with dexamethasone (2–3 mg kg−1) and analgesics (carprofen; 5 mg kg−1). A subgroup of animals was also injected with atropine (0.05–0.1 mg kg−1). General anaesthesia was induced either with a mixture of fentanyl (0.05 mg kg−1), midazolam (5 mg kg−1) and medetomidine (0.5 mg.kg−1), or with isoflurane (1%–4%). A custom headplate was attached to the skull using dental cement (C&B Super Bond) and the skull above the posterior cortex was carefully thinned and sealed with a thin layer of light-cured dental composite (Tetric EvoFlow). Viral injections of AAV1.EF1a.DIO.hChR2(H134R)-eYFP.WPRE.hGH (Addgene20298) (titre: 2.2 × 10^12^, diluted 1:4 in cortex buffer (described below), 70 nl) were made using glass pipettes and a pressure injection system (Picospritzer III, Parker) in the right hemisphere in either V1 or LM. The injection site was identified using intrinsic imaging maps of visual cortical areas (see Intrinsic signal imaging) several days prior to the surgery. Some animals were additionally given antibiotic and analgesic drugs (enrofloxacin 5 mg kg−1, buprenorphine 0.1 mg kg−1) at the end of surgery and for 3 days during recovery. Electrophysiological recordings were performed approximately 2-4 weeks after the viral injections.

### Intrinsic imaging

To determine retinotopically matched locations in V1 and LM, mice underwent optical imaging of intrinsic signals^69^. This procedure was done a minimum of 3 days after the implantation of a headplate and thinning of the skull (see surgical procedures). On the day of intrinsic imaging, mice were either initially sedated (chlorprothixene, 0.7 mg kg−1), and imaging was carried out either under light isoflurane anaesthesia (0.5–1%) delivered via a nose cone, or imaging was performed in awake, head-fixed mice, free to move on a 20-cm-diameter Styrofoam cylinder.

The visual cortex was illuminated with 700-nm light split from a LED source into two light guides. Imaging was performed with a tandem lens macroscope focused 500 µm below the cortical surface and a bandpass filter centred at 700 nm with 10 nm bandwidth (67905; Edmund Optics). Images were acquired with a rate of 6.25 Hz with a 12-bit CCD camera (1300QF; VDS Vosskühler), a frame grabber (PCI-1422; National Instruments) and custom software written in Labview (National Instruments). The visual stimulus, presented on a display 21 cm away from the left eye, was generated using the open-source Psychophysics Toolbox^70^ based on Matlab (MathWorks) and consisted of a 20° (radius) large square-wave grating, (0.08 cycles/degrees) drifting at 4 Hz in 8 random directions, presented on a grey background for 2 seconds, with a 18 second interstimulus interval alternatively at two positions, at 15° elevation and either 50° or 70° azimuth. Frames in the second following stimulus onset were averaged across 16 grating presentations to generate intrinsic response maps. Response maps to the grating patches at either position were used to identify retinotopically matched locations across areas V1 and LM.

### Visual discrimination task and visual stimuli

Mice were trained for 2 - 8 weeks to perform a Go/No-go task, in which they had to discriminate two static gratings of 45° and -45°orientation. Mice were food restricted throughout the training and electrophysiological recording, with maximum weight loss of 20% of their initial body weight. The restriction started 3 days after the first surgery (headplate implantation) and was interrupted for several days after viral injections, for mice to recover after surgery. The mice were trained for the duration of approximately 1 hour every day. Initially, mice were trained to run head-fixed on a freely rotating Styrofoam cylinder in front of a display (Dell U2715H, 60Hz) placed 24 cm away from their left eye. In later stages of training, when mice were comfortable and used to the paradigm, the cylinder was fixed in place, preventing it from moving. Mice learned within a few days to transition from running to sitting still while performing the task.

Visual stimuli consisted of two 40° diameter large static square-wave gratings of 100% contrast, 0.06 cycles per degree spatial frequency, and either 45° or -45° orientation, presented on a grey background in the centre of the monitor. The centre of the monitor was placed at approximately 60° azimuth and 15° elevation, and was adjusted later according to the receptive fields of recorded neurons (described below). The luminance of the monitor was 0 *cd*/*m*^2^, 16 *cd*/*m*^2^, and 32 *cd*/*m*^2^ at black, grey and white values, respectively. Gratings were presented for 500ms. The interstimulus interval consisted of a fixed period of 500 ms, plus a random delay with a truncated exponential distribution to avoid extremely long trials (mean: 4-6 seconds, truncation threshold: between 10 and 30 seconds).

6 mice were trained with the 45° grating as the Go and the -45° stimulus as the No-go stimulus. For the remaining 10 animals, the identity of Go and No-go stimuli was switched. Go and No-go stimuli occurred with equal probability. We did not find any stimulus-specific differences in V1 and LM activity or their interactions, therefore data was pooled across all animals. A reward delivery spout was positioned under the snout of the mice from which a drop of soy milk, or Ensure Plus strawberry drink was delivered in Go trials triggered by licking of the spout during a response window of 100 to 550ms (n = 8 animals) or 100 to 650ms (n = 8 animals) after stimulus onset. If mice licked the spout during this time window in response to the Go stimulus, trials were classified as hit trials, otherwise as miss trials. In the miss trials, mice received an automatic reward (smaller drop of reward) after the response window. The same time window was used to classify No-go trials into false alarm or correct rejection trials. Licking to the No-go visual stimulus (false alarm) was not punished. Licks were detected with a piezo disc sensor placed under the spout. The detection of licks, reward delivery, and the presentation of visual stimuli were controlled by a MATLAB-based script, StimServer^71^ (an open-source stimulus sequencing package based on the Psychophysics Toolbox), and using a data acquisition board (PCIe-6321; National Instruments).

### *In vivo* electrophysiology and optogenetic manipulations

Electrophysiological recordings were performed 2 - 4 weeks after the viral injection. Most mice were trained in the visual discrimination task prior to the recording day. On the day of the recording, mice were anaesthetized with 1%–2% isoflurane, and two 1 mm craniotomies were made above the pre-selected retinotopically matched locations in V1 and LM (see intrinsic imaging). Craniotomies were covered with 1.5-2% agarose in cortex buffer, containing (in mM) 125 NaCl, 5 KCl, 10 Glucose monohydrate, 10 HEPES, 2 MgSO4 heptahydrate, 2 CaCl2 adjusted to pH 7.4 with NaOH. A well was built around the two craniotomies, using light-cured dental composite (Tetric EvoFlow), and finally, the well was further sealed with Kwik-Cast sealant (World Precision Instruments).

Mice recovered from surgery for 1-2 h before the recording, and were then head-fixed on a styrofoam cylinder, free to move prior to the start of recording. The Kwik-Cast sealant was removed and a silver wire was placed in the bath for referencing. Two NeuroNexus silicon probes (A2x32-5mm-25-200-177-A64), labelled with DiI, were lowered to 800-1000 μm below the cortical surface using micromanipulators (Sensapex). The electrode positions in V1 and LM were chosen based on the intrinsic map (see intrinsic imaging), to target retinotopically matched and therefore anatomically connected locations in the two areas^54,72^ (**Extended Data Fig. 1a**). The electrode positioned in the injection site (in V1 for V1 silencing and in LM for LM silencing) had a 200*μm* optical fibre (CFMLC52U; Thorlabs) attached with dental cement with its tip 1mm above the electrode tip, placing the fiber tip on the cortical surface once the electrodes were positioned in the cortex.

The craniotomies were then covered with 1.5-2% agarose in cortex buffer. Voltages from 128 channels across two areas were acquired through amplifier boards (RHD2132, Intan Technologies) at 30 kHz per channel, serially digitized and sent to an Open Ephys acquisition board^73^ via a SPI interface cable. The wheel was fixed in place (no movement possible), and mice sat quietly on the wheel due to the prior training. Before the start of the behavioural paradigm, retinotopic mapping of the neurons in V1 and LM electrodes was performed, using flashing black and white squares of approximately 20° × 20° on a grey background in 5×4 locations across the monitor. Responses to these stimuli were used to position the monitor, centred on the average receptive field centre of recorded neurons.

To silence neuronal activity in either V1 or LM, we optogenetically activated ChR2-expressing parvalbumin-positive neurons using a 473 nm laser (OBIS 473nm LX 75mW; Coherent) or LED (M470F3; Thorlabs) coupled through a patch cable (M73L01; Thorlabs) to the fibre above the area previously injected with the AAV. The light pulse lasted for 150ms. During the first 75 ms, the power was constant at 4 mW, and during the second 75ms, the power was linearly ramped down to zero, in order to prevent rebound spiking. The light pulse was generated using Pulse Pal (open source pulse generator). Optogenetic silencing effectively suppressed neural activity (Median change in the firing rate of V1 neurons was -95.74% when silencing V1, and -93.02% for LM neurons when silencing LM). Propagation of light to the eye was blocked by the cement wall around the craniotomies, as well as black tape shielding the fibre-mating sleeves. Since the fibre tip was far from the eye (> 5 mm) light did not propagate to the retina through the brain^74^. Light was delivered during 8 different time windows during the presentation of the visual stimulus. The onset of light delivery with respect to visual stimulus onset (silencing onset) was randomized from trial to trial. The exact alignment of light onset and visual stimulus was confirmed offline, using recordings of the input pulse to the laser, and the monitor frame update times captured by a photodiode attached to the monitor. These onsets were at 0 ms, 56ms, 123ms, 189ms, 256 ms, 323ms, 390ms and 456 ms after visual stimulus onset (mean across n = 14 animals). We ensured minimal trial-to-trial jitter in laser onset delays (standard deviation of light onset time < 0.3 ms). Control trials without light were interspersed randomly. One session was recorded per mouse, and each session consisted of on average 78 ± 8 (mean ± s.d.) silencing trials for each time window, and 97 ± 16 (mean ± s.d.) control trials (before excluding trials based on the criteria described below).

## Data Analysis

### Analysis of behaviour

A behavioural d-prime was calculated for each animal (d’ = Z(hit rate) - Z(false alarm rate), where function Z is the inverse of the standard normal cumulative distribution function), and only animals with task performance above chance level were included for further analysis (2 animals out of 16 excluded). Chance level was the 99 percentile of the trial-shuffled d-prime distribution, obtained by shuffling the identity of Go and No-go trials 5000 times. Average task performance for the animals included in the analysis was d’ = 1.7 ± 0.16 (mean ± s.e.m). Only correct trials (hit and correct rejection trials) were used for subsequent analyses. Moreover, periods of time in the session during which the mice were grooming (continuous prolonged movement detected from the spout readout) were excluded.

### Electrophysiology data pre-processing

Spikes were sorted with Kilosort (https://github.com/cortex-lab/Kilosort) using procedures previously described^75^, single units were extracted, and manually curated using phy^76^. A total of 806 single units in the trained animals (n = 14 mice, 371 units in V1 and 435 units in LM) and 236 single units in untrained animals (n = 4 mice, 146 units in V1 and 90 units in LM) were detected. For all subsequent analysis, the firing rate in time windows with an average firing rate below a threshold was set to nan. This was done in order to prevent dividing by values close to zero. The threshold was chosen as the value at which the 95% confidence interval of the firing rate exceeded zero (2.5 Hz). Results were not significantly different when neuronal activity was not thresholded.

### Receptive field calculation

Receptive fields were calculated by fitting a two-dimensional Gaussian distribution to the responses of neurons to the flashing black and white square stimuli. The centre of the receptive field was calculated as the peak of the Gaussian fit. Since receptive fields were calculated from square stimuli on a flat screen, the receptive field locations in visual degrees are approximate. However, this does not affect the relative comparisons between the regression models in V1 and LM (described below).

### Quantifying the effect of silencing

The change caused by optogenetic manipulation to the firing rates of individual neurons was quantified as 100 × (*R_s_* − *R_c_*)/*R_c_* for each silencing time window, where *R_s_* denotes the cell’s spike count during the 150ms silencing time window, and *R_c_* denotes the spike count in the corresponding time window in control trials. Significant effects were determined based on exceeding the 95% confidence interval of percentage differences in the firing rate of bootstrapped control trials. To determine the number of significantly affected time windows per cell, the significance levels were adjusted using Bonferroni correction for 8 comparisons.

### Multivariate regression model

We used a generalized linear regression model in order to examine if silencing effects could be explained by physiological and anatomical properties of target neurons such as cortical depth. We predicted the activity of LM neurons (target area) during optogenetic silencing of V1 (source), from their own activity in control trials (in the corresponding time bin) and the depth of LM neurons in the cortex (**Extended Data Fig. 2**):

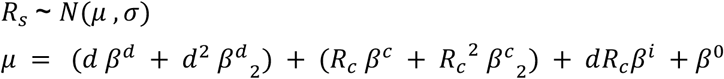

Where *R_s_* denotes the spike count during the 150 ms source-silencing time window, modeled with a Gaussian distribution with the mean and standard deviation of *μ* and *σ* . *R_c_* denotes the spike count in the corresponding window in control trials, *d* the depth of the target neuron soma, and *β* the model coefficients. Cortical depth was determined based on current source density analysis (CSD), with zero denoting the centre of the initial current sink (which roughly corresponds to thalamic input in layer 4 in V1). The model was fit with the iteratively reweighted least squares algorithm, using the fitglm function from MATLAB. The performance of the model was assessed using 20-fold cross validation. For each set of training and test sets, the model parameters were fit using the training set, and the log likelihood of the test set was calculated (*log*_2_ *l*). We also used the same procedure to calculate the log likelihood of a null model that only contained information about the LM cell’s control trial activity (*μ*_null_ = *R_c_ β*^*c*^ + *R_c_*^*c*^*β*^*c*^_2_ + *β*^0^). This gives a null likelihood value (*log*_2_*l*_0_).

We then measured model performance as the degree to which the full model outperforms the null model by calculating the trial-averaged excess log likelihood of the full model compared to the null model. This would capture any dependence of the silencing effect on the depth of LM neurons, that is not solely due to the relationship between activity levels and cortical depth.

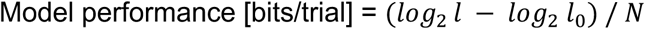

Where *N* is the number of observations in the test dataset. Model performance was calculated for each of the 20 test sets (observations) separately. Above-chance performance was determined by a one-sided Wilcoxon signed-rank test. We constructed similar, separate models to quantify the influence of other cell properties, such as relative receptive field position and average firing rate over the session. The relative receptive field positions were the distance of the cell’s spatial receptive field centre to the retinotopic location of silencing in the source area, which was calculated as the average receptive field location of the recorded neurons from an electrode positioned in the centre of AAV-flex-ChR2 virus expression in the source area. We repeated the same procedures for predicting the activity of V1 neurons during optogenetic silencing of LM.

### Causal Communication Direction

We characterized the influence of the source area on target neurons on the population level. For a population of *n* simultaneously recorded neurons in the target area, we found an *n* × 1 vector, in the *n* dimensional activity space that maximally separated the activity vectors in the control and source-silencing trials. This was done separately for each silencing time window and stimulus type (Go/ No-go). The source-silencing activity vectors corresponded to the population firing rate of the n neurons in the 150 ms in which the source area was optogenetically silenced, and the control activity vectors were the firing rate of the same population in the corresponding 150 ms time window in control trials. In order to find the direction maximally separating the control and source-silencing trials, we used regularized linear discriminant analysis (LDA). Communication direction (*CD*) was defined as the unit-length normal vector to the decision boundary hyperplane.

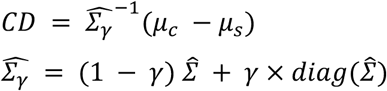

Where 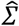 denotes the pooled covariance estimate, *μ_c_*, *μ_s_* correspond to the mean activity vectors, and *γ* is the regularization parameter. In addition to *γ*, we used another parameter for each classifier. Any coefficients in *CD* with a magnitude smaller than a threshold, *δ*, was set to zero, eliminating the corresponding neuron from the discriminant direction.

The two parameters of discriminant classifiers (*γ*, *δ*) were fit separately for each animal by minimizing the test error in a ten-fold cross validation. Excluding *δ* did not change results significantly. The classifier performance for the LDA classifiers in trained animals are shown in **Extended Data Fig. 4a-f**. All included animals had above chance classification performance.

For the above analysis, only animals with at least 10 control trials (n=7 of 7 and n=6 of 7 mice for V1 and LM silencing, respectively), and only visually responsive cells were included. Neurons had to spike at least once (on average) during the 500 ms stimulus duration in response to the Go or No-go stimulus, and there needed to be a significant difference (p = 0.05, two-sided Wilcoxon signed-rank test) between the spike count during the stimulus and a 500 ms window before stimulus onset for at least one of the stimuli. An equal number of Go and No-go trials were used for calculating the communication direction at each time window.

The similarity of two directions (cosine angle) was captured by their dot product *cos(α_i,j_)* = (*CD_i_*.*CD_j_*)/(|*CD*_*i*_|. |*CD*_*j*_|), represented in the *i*th row and *j*th column of the cosine similarity matrix. In order for this measure to reflect trial-to-trial stability of the communication directions, we used a cross-validated similarity measure. We randomly split the control trials in half, one half was used to calculate *CD_i_* and the other, *CD_j_*. This was then repeated in reverse, and the resulting dot products were averaged. The random partitioning of trials was done 100 times, and the average dot products were reported in the cosine similarity matrices. This method accounts for trial-to-trial variability and summarizes robustness of cosine similarity to trial-to-trial noise. Deviation from 1 of the diagonal elements in the cosine similarity matrix (cosine similarity at lag 0) reflect the trial to trial variability of communication directions at each time window.

Comparing differences in average activity vectors (not normalized to covariance), instead of regularized LDA vectors gave similar results (**Extended Data Fig. 5g-j**). However, we used regularized LDA in order to find the mode of activity which would most reliably specify the difference between control and silencing trials across trials.

In order to track the similarity of trial-averaged population activity over time, we defined activity directions at each time window as the *n* × 1 unit length vector specifying the coordinates of trial-average population activity. To calculate the activity directions, we used 65 ms time windows (instead of the full 150 ms time windows used elsewhere), in order to avoid temporal overlap between successive time points. Using 150 ms windows for activity directions resulted in even slower changes over time (data not shown). The activity directions were calculated from V1 and LM activity in control trials, pooled across all animals included in the LM silencing or V1 silencing experiments. The similarity of activity directions was assessed using cross-validated cosine similarity as explained above. Similarity in activity directions implies that different neurons undergo similar temporal fluctuations of activity, leading to a change only in the magnitude of population activity, and not its direction over time (**Extended Data Fig. 1c**). For comparing how fast similarity of activity and communication directions decay over time, we used the initial slope (slopes between lag 0 and lag 1) of the decay over time lags of pairwise similarities, (decay slope of similarity between adjacent time windows). This initial decay slope, as opposed to exponential decay time constant, does not assume an underlying exponential decay, and can be used as a general measure. We chose this measure since the similarity of activity directions decays linearly rather than exponentially (different to the similarity of communication directions) over time.

### Sub-selection of cells in the source area

To rule out the possibility that the fast changes in the similarity of communication directions over time arise from changes in the activity direction of a select subnetwork of neurons in the source area (e.g., projection neurons) with particularly fast dynamics, we examined the activity directions of the subgroup of source neurons with the most dynamic firing rate compared to the average population. In order to identify a subpopulation of neurons in the source area with the fastest decay in their similarity of activity vectors, we z-scored the activity of all simultaneously recorded source neurons at each time window separately. Then, we selected 20% of cells with the highest standard deviation over time in their z-scored activity. This procedure leads to selecting subpopulations that have the highest deviation from the average population activity over time, and would therefore have the fastest change in their activity vector similarity over time. In order to compare the cosine similarity of activity directions derived from this subpopulation to those of the communication directions, we re-calculated the communication directions from a randomly selected subpopulation of 20% of target neurons (averaged over 100 repeats), This sub-selection of target neurons was done in order to make the maximum dimensionality of communication directions comparable to those of the source area activity directions.

### Simulation of time invariant inter-areal influences

In order to simulate time-invariant influence of areas on each other, we generated a dataset in which the activity of each target neuron during source-silencing was simulated based on its control activity and assuming similar source-silencing effects during all time windows. For this, for each target neuron, the source-silencing effect of one of the 8 silencing time windows (*E^T^*) was selected randomly and used for all time windows as the cell’s time-invariant silencing effect (*E*), with the randomization repeated 100 times. Next, for each silencing time-window, the source-silencing spike count was simulated by first generating a Poisson sample with the mean of control trial spike counts in the same time window (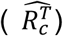), and then, subjecting the activity of each simulated trial to the time-invariant silencing effect (*E*), to get the simulated source-silencing spike counts (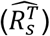):

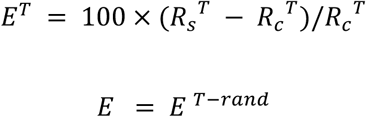

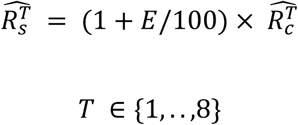

Where *R_s_*^2^ and *R_c_*^2^ are the spike count in source-silencing and control trials during the silencing time window T. By generating spike counts using a Poisson distribution, the trial-to-trial variability of target neuron activity is preserved, while the underlying source-silencing effect is assumed to be time-invariant. Since the time-invariant source-silencing effect on each target neuron is selected randomly based on its own effects, over 100 repeats, the time-averaged source-silencing effects are similar to the experimental dataset, and the overall effect size of source-silencing was preserved in the simulated dataset. Importantly, since the simulations are based on the control firing rate of target neurons in each time window which contain input from the source area, we account for the compound trial-to-trial variability in target area activity and source area input.

### Principal component analysis

The principal components (PCs) of V1 activity were calculated separately for the population activity during each 150 ms time window, either in the control or LM silencing trials. PCs were calculated in each time window separately for Go and No-go trials, and for control and silencing conditions, with an equal number of trials used across the 4 conditions. In order to calculate the similarity of PC directions over time, we used the absolute value of cross-validated similarity as described above (see causal communication direction section), where only half of the trials were used to calculate PCs at each time window. To calculate the total variance and eigenspectra, we used all included trials at each time window to calculate the variance along each principal component. Similarly to the communication direction analysis, only animals with at least 10 control trials, and only visually responsive cells were included in the analysis (for responsiveness criteria see causal communication direction section). To quantify the decay of PC similarity across time lags without explicitly assuming an exponential decay, we used the decay slope described above (see causal communication direction section).

### Exponential fits

For comparing time constants of exponentially decaying functions, the decay of cosine similarity over time lags was fit by an exponential decay with an offset:

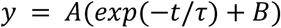

Where *t* is the time lag between two silencing onset times. Data was fit using nonlinear least-squares fitting via the trust-region algorithm (through MATLAB fit function). Confidence intervals of the fit parameters were obtained using a 100-times bootstrap. To assess the significance of the differences between the decay time constants of communication during Go and No-go stimuli, we used a permutation test. To do this, we calculated the distribution of time constant differences under the null hypothesis. This null distribution was generated from exponential fits on data with Go and No-go trial labels exchanged randomly (1000 randomized permutations).

### Autocorrelations and pairwise noise correlations

To calculate the temporal autocorrelation of spike counts, we divided the stimulus duration into successive time bins of 65 ms duration. We then calculated across-trial Pearson’s correlation coefficient between the spike counts at any two time bins and plotted against the time lag between the two time bins. This was done separately for each neuron. This method subtracts the across-trial mean spike counts, therefore the spike-count autocorrelations don’t explicitly depend on mean firing rates at different time bins.

Noise-correlation strength was calculated for each pair of neurons in V1 as the across-trial Pearson’s correlation coefficient between the spike counts of the two neurons at a given time bin, using similar time bins as for the autocorrelation analysis. These correlation coefficients were plotted as a function of the average spike count of the pair in the same time bin and values calculated from different time bins were then pooled. For calculating autocorrelations and pairwise noise correlations, similarly to the communication direction analysis, only animals with at least 10 control trials, and only visually responsive cells were included in the analysis (for responsiveness criteria see causal communication direction section), and an equal number of trials was used across Go and No-go trials.

## Acknowledgements

We thank Thomas Mrsic-Flogel, Ryan Low, Grace Lindsay, Ivan Voitov, Joana Soldado-Magraner, and Antonin Blot for comments on the manuscript. We thank Michelle Li for animal husbandry. We thank Thomas Mrsic-Flogel, Maneesh Sahani, Peter Latham, Kenneth Harris, Maxime Rio, Antonin Blot, and Ivan Voitov for feedback and discussions. This work was supported by the Sainsbury Wellcome Centre Core Grant from the Gatsby Charitable Foundation and Wellcome (090843/F/09/Z), by an ERC Starting Grant (SBH, HigherVision 337797) and by Biozentrum, University of Basel core funds.

## Author Contributions

M.J. and S.B.H. conceptualized the study. M.J. performed experiments and analysed the data. S.B.H. and M.J. wrote the manuscript.

Correspondence to Mitra Javadzadeh or Sonja B. Hofer.

## Data and materials availability

Data, software used for behavioral training, and analysis code is available from the corresponding authors upon reasonable request.

---

**Extended Data Fig. 1.**
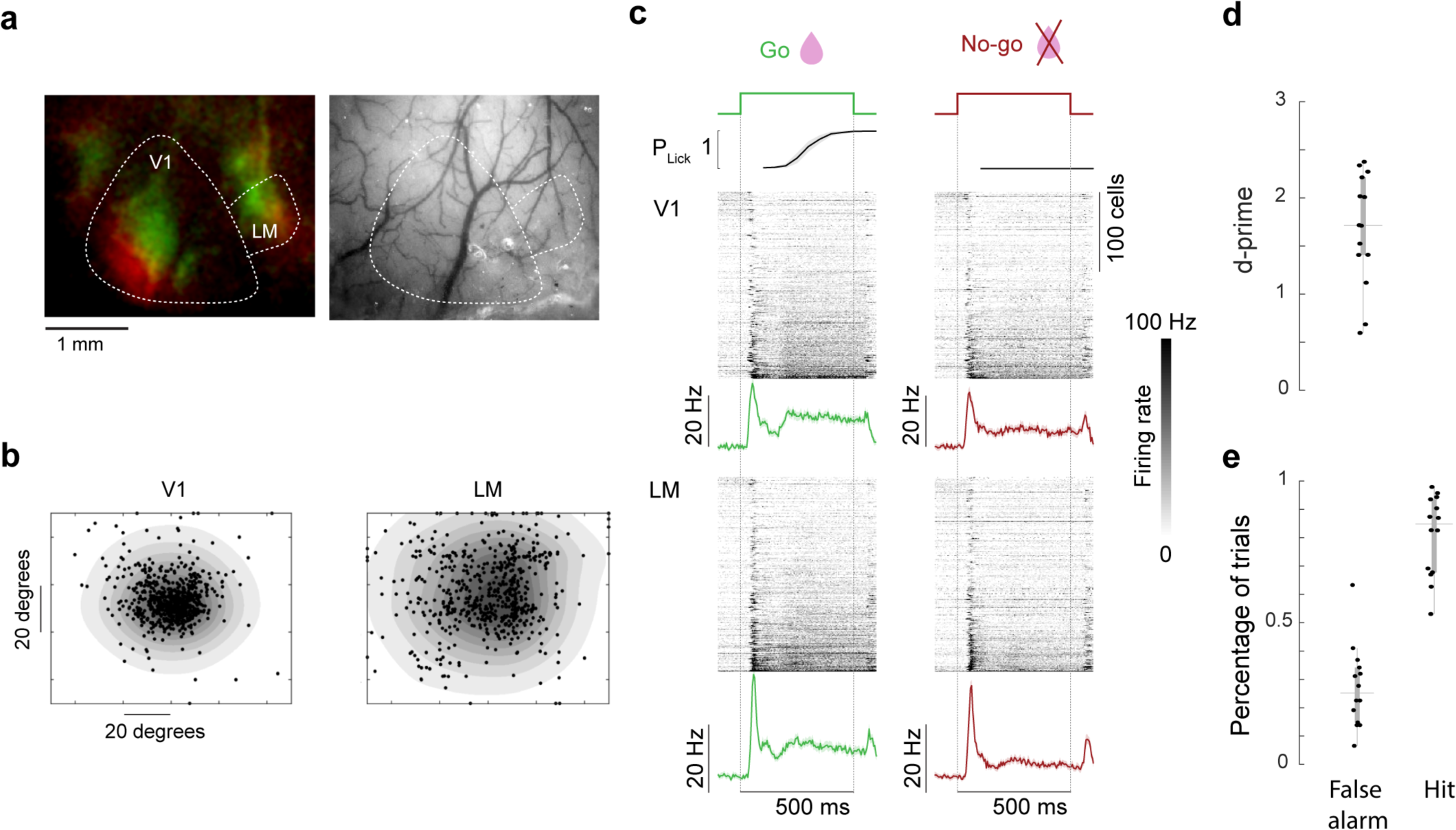
Targeting retinotopically matched regions in V1 and LM using intrinsic signal imaging. **a,** Left: example intrinsic imaging map used to target V1 and higher visual lateromedial area (LM). Intrinsic responses evoked by two spatially separated visual stimuli (see Methods) are colour-coded in green and red. Right: surface blood vessel pattern corresponding to the intrinsic imaging map on the left. Approximate borders of V1 and LM are outlined. **b,** Receptive field centres of recorded neurons in V1 (left) and LM (right) (n = 16 animals) overlaid on the average receptive field of all neurons, obtained by individual 2D Gaussian fits. **c,** Top, cumulative lick probability over time from stimulus onset (averaged over trials and mice, n = 14 mice) in correct Go (left) and correct No-go (right) trials. Shading depicts 95% confidence interval of the mean. Middle and bottom, spiking activity of neurons in V1 (middle) and LM (bottom) in response to the Go (green, left), and the No-go (red, right) stimulus. Each row in the heat plots denotes the average firing rate of one neuron. The traces below are average PSTHs of all recorded visually responsive neurons, binned at 20 ms. **d,** Task performance (behavioural d-prime, see Methods) of all animals included in the analyses (n = 14). The criterium for inclusion in the analyses was task performance above chance level, calculated as the 99 percentile of the trial-shuffled d-prime distribution (see Methods). **e,** Percentage of false alarm and hit trials for all animals included in analyses (n = 14).

**Extended Data Fig. 2.**
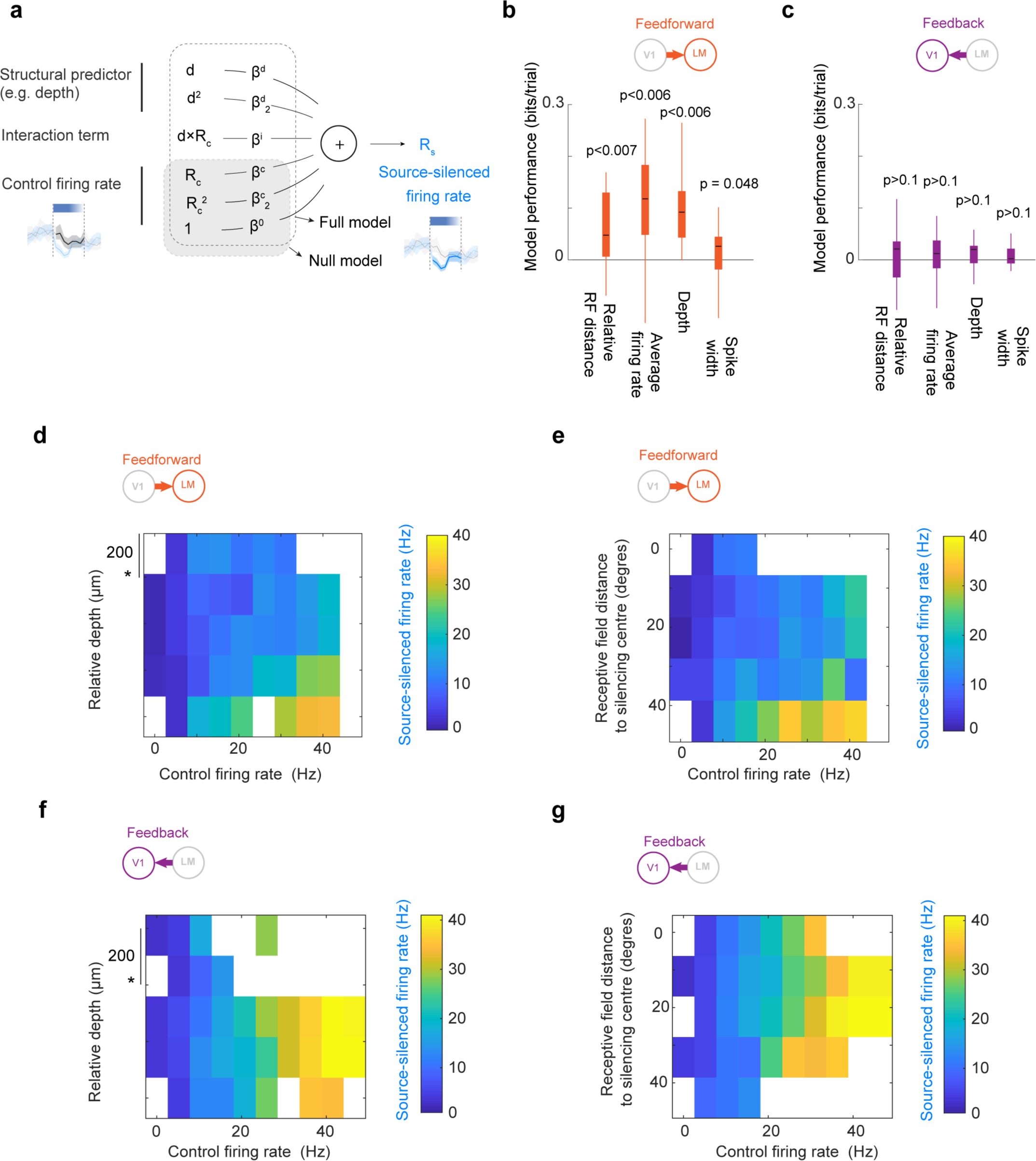
Dependence of firing rates during source silencing on the properties of target neurons. **a,** Regression model predicting the activity of target neurons during optogenetic silencing of the source area, from a combination of the neurons’ activity in control trials in the same time window and the structural predictor of interest (e.g. cortical depth). The null model contains information only about the control trial activity. Models were fit separately for the following predictors: distance of spatial receptive field centre from the retinotopic location of the centre of optogenetic silencing in the source area, average firing rate during the session, depth of neuron in the cortex, and the width of action potential waveforms. **b,** The cross-validated performance of each model predicting the influence of V1 silencing on LM cells (feedforward influence), as the degree to which the model outperforms the null model (in bits/trials). Y axis indicates the log likelihood ratio (full model to null model) of the test observations in each cross-validation fold (n=20), normalized to the number of observations. These values would capture any dependence of the influence of V1 on LM neurons’ activity on the particular predictor (indicated on the x axis) that is not explained by the modulation of firing rates by the predictor in control conditions. Values significantly above zero (p-value < 0.05, Bonferroni correction, one-sided Wilcoxon signed-rank test) indicate above-chance prediction power of the model. **c,** As in **b,** but for predicting the firing rate of V1 neurons during LM silencing. **d,** Firing rate of LM neurons during V1 silencing (indicated by colour code) as a function of their firing rate in control conditions without optogenetic manipulation (control firing rate, x axis) and their depth in the cortex (y axis). Modulation in the vertical axis (for a given control firing rate) indicates the effect of depth on the neurons’ activity in the absence of V1 input. The star (*) denotes the centre of the initial current sink detected from current source density analysis (in V1 this corresponds approximately to layer 4). **e,** Firing rate of LM neurons during V1 silencing as a function of their control firing rate (x axis) and the distance of their receptive field centre from the retinotopic location of the centre of optogenetic manipulation in V1 (y axis). Modulation in the vertical axis (for a given control firing rate) indicates the effect of relative receptive field position on the neurons’ activity in the absence of V1 input. **f,** as in **d,** but for the firing rate of V1 neurons during LM silencing. **g,** as in **e,** but for the firing rate of V1 neurons during LM silencing.

**Extended Data Fig. 3.**
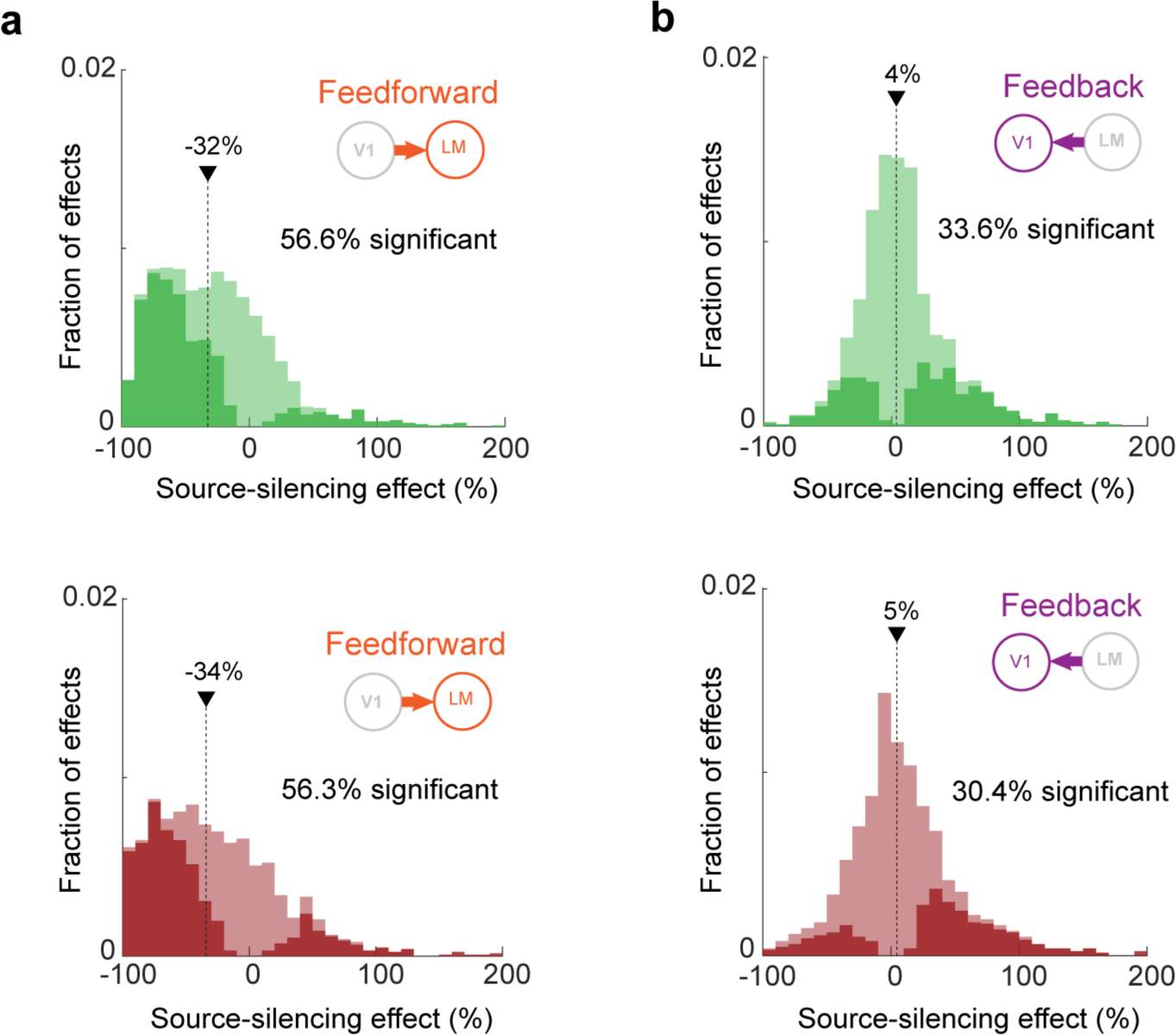
– Distribution of silencing effects in Go and No-go trials. **a,** Distribution of the effect of silencing V1 on individual LM neurons (feedforward influence) in all silencing time windows during Go trials (green, top) and No-go trials (red, bottom). Significant effects shown in bright colours. The arrow denotes the median of the distribution. Go vs No-go p = 0.87, two-sided Wilcoxon rank-sum test. **b,** Distribution of the effect of silencing LM on individual V1 neurons (feedback influence) in all silencing time windows during Go trials (green, top) and No-go trials (red, bottom). Significant effects shown in bright colours. The arrow denotes the median of the distribution. Go vs No-go p = 0.78, two-sided Wilcoxon rank-sum test.

**Extended Data Fig. 4.**
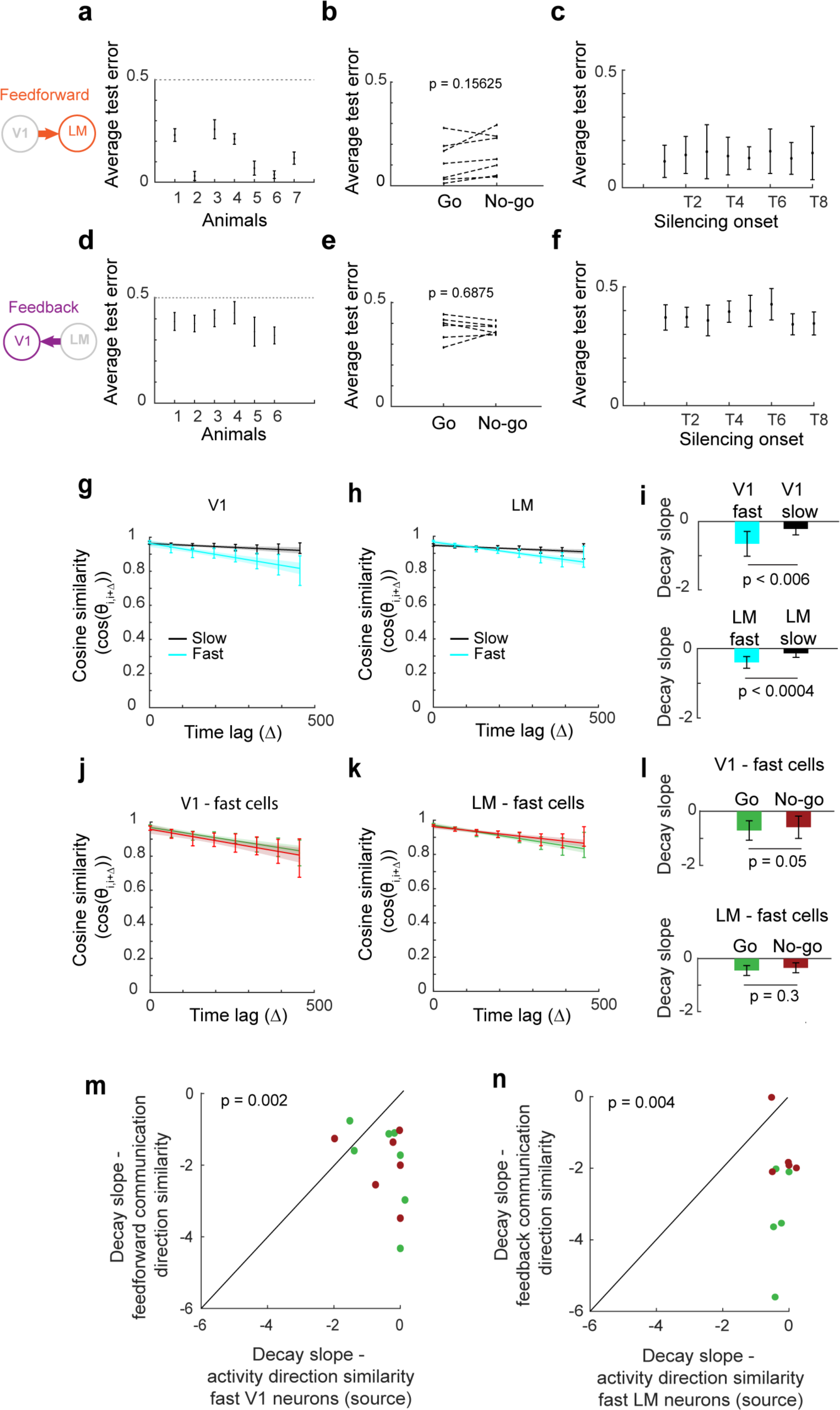
-LDA and communication direction controls. **a,** Cross-validated performance of the regularized linear discriminant analysis (LDA) for distinguishing LM population activity in control trials from that in V1 silencing trials. Y axis shows the percentage of misclassified trials (pooled across Go and No-go stimuli) in the test data set using 10-fold cross validation for each individual animal. **b,** Cross-validated performance of the regularized linear discriminant analysis for distinguishing LM population activity in control trials from that in V1 silencing trials during Go and No-go stimuli (p = 0.156, two-sided Wilcoxon signed-rank test). Points connected by dotted lines indicate individual animals. **c,** Cross-validated performance of the regularized linear discriminant analysis for distinguishing LM population activity in control trials from that in V1 silencing trials in the eight silencing time windows with varying onset times, averaged across animals (N = 7 animals). Performance was similar for the eight silencing time windows (one-way ANOVA p = 0.99). **d,** as in **a,** but for distinguishing V1 population activity in control trials from that in LM silencing trials. **e,** as in **b,** but for distinguishing V1 population activity in control trials from that in LM silencing trials. **f,** as in **c,** but for distinguishing V1 population activity in control trials from that in LM silencing trials. (N = 6 animals, one-way ANOVA p = 0.37). **g,** Pair-wise cross-validated cosine similarity of activity directions in different time windows as a function of the time lag between them in two subpopulations of V1 neurons, the 20% of neurons with the most time-varying activity (based on the standard deviation over time in their z-scored activity, ‘fast’, blue), and the 20% of neurons with the least time-varying activity (‘slow’, black). Data was pooled across Go and No-go stimuli (n = 13 mice). **h,** as in **g,** but for LM cells. **i,** Initial slope (slope between lag 0 and lag 1) of the decay over time lags of fast-changing and slow-changing V1 subpopulations in **g,h**. Error bars depict the 95% confidence interval of the mean (2×s.e.m). **j,** Pair-wise cross-validated cosine similarity of activity directions in different time windows as a function of the time lag between them in fast-changing V1 cells (20% of V1 neurons with the most time-varying activity) during Go (green) and No-go trials (red). **k,** As in **j,** but for LM cells. **l,** Top, initial slopes (slopes between lag 0 and lag 1) of the decay over time lags of fast-changing V1 neurons (20% of V1 neurons with the most time-varying activity) in Go (green) and No-go (red) trials. Error bars depict the 95% confidence interval of the mean (2×s.e.m). Bottom, as on top, but for the fast-changing LM neurons. **m,** Relationship between the initial slopes (slopes between lag 0 and lag 1) of the decay over time lags of feedforward communication direction similarities and of activity direction similarities in fast-changing V1 neurons (20% of V1 neurons with the most time-varying activity, see Methods) in Go (green) and No-go (red) trials for individual animals (n = 7 animals). In order to make the maximum dimensionality of communication directions comparable to those of the source area activity directions, the communication directions were re-calculated from a randomly selected subpopulation of 20% of target neurons (see Methods). **n,** Relationship between the initial slopes (slopes between lag 0 and lag 1) of the decay over time lags of feedback communication direction similarities and of activity direction similarities in fast-changing LM neurons (20% of LM neurons with the most time-varying activity, see Methods) in Go (green) and No-go (red) trials for individual animals (n = 6 animals). In order to make the maximum dimensionality of communication directions comparable to those of the source area activity directions, the communication directions were re-calculated from a randomly selected subpopulation of 20% of target neurons (see Methods).

**Extended Data Fig. 5.**
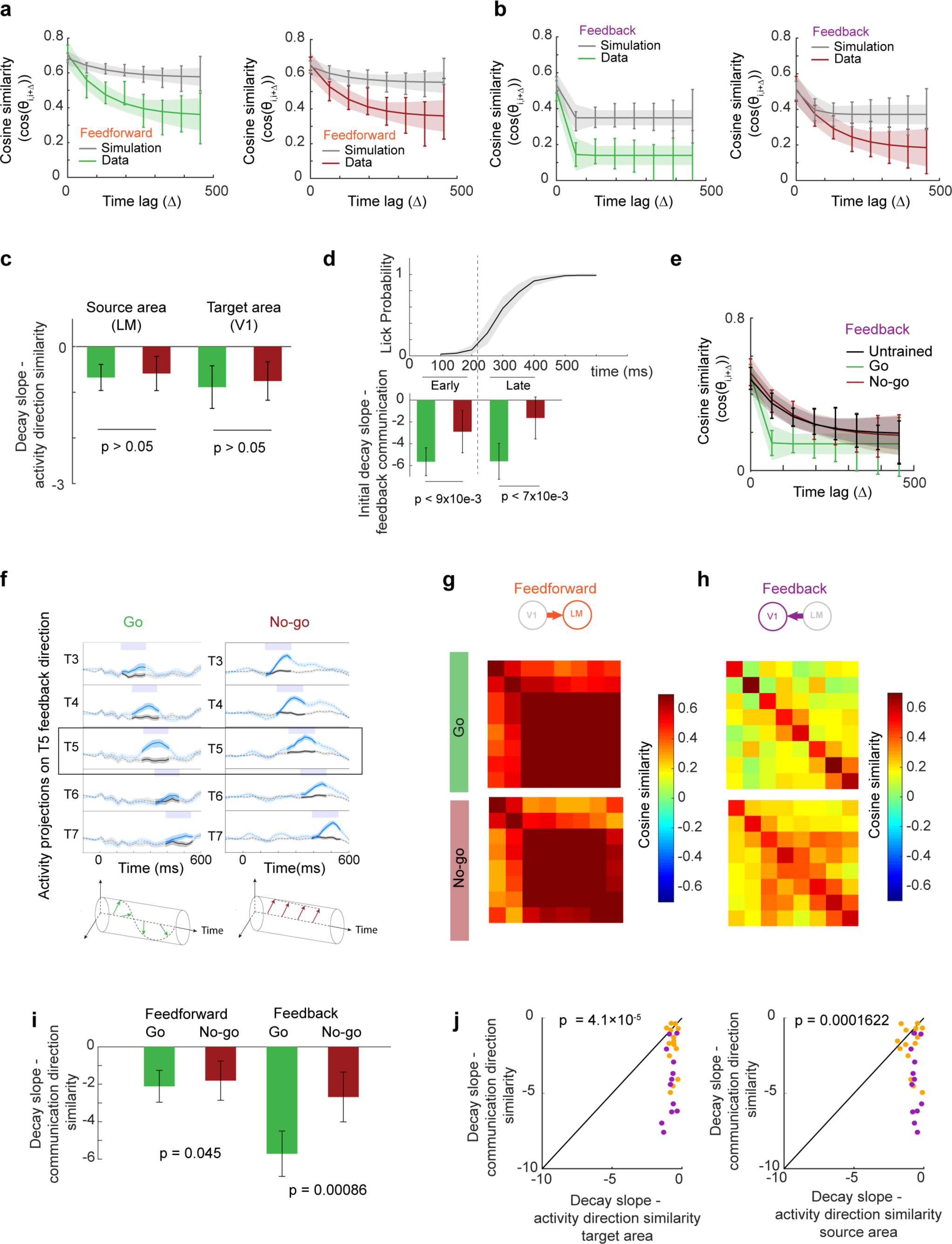
-Communication direction and activity direction controls. **a,** Cross-validated cosine similarity of pairs of feedforward communication directions (influence of V1 silencing on LM activity) in different time windows as a function of the time lag between them during the Go (left, green) and No-go stimulus (right, red) and the corresponding cross-validated cosine similarity functions from the simulated dataset with time-invariant feedforward influences (grey). Error bars depict the 95% confidence interval of the mean (2×s.e.m). Lines depict exponential fits, shading shows 95% prediction bounds of the fits. **b,** As in **a,** but for feedback communication directions (influence of LM silencing on V1 activity) **c,** Initial slope (slope between lag 0 and lag 1) of the decay over time lags between activity directions during Go (green) and No-go (red) trials, describing population activity in area LM (left) and V1 (right) in experiments in which LM silencing was performed (n = 6 animals). Error bars depict the 95% confidence interval of the mean (2×s.e.m). **d,** Top, cumulative lick probability in correct Go trials over time from stimulus onset (averaged over trials and mice). Bottom, initial decay slope of communication direction similarity (between time lags 0 and lag 1, similar to Fig. 3l), for LM feedback influences on V1 activity, measured early during visual stimulus presentation (before most licks, silencing onsets 56, 123 and 189 ms after stimulus onset) and in a late period (during licking, 256, 323 and 390 ms after stimulus onset). Error bars depict the 95% confidence interval of the mean (2×s.e.m). **e,** Cross-validated cosine similarity of pairs of feedback communication directions in different time windows as a function of the time lag between them during grating stimuli presentation in untrained animals (black, passively viewing the visual stimuli), compared to trained animals during Go (green) and No-go (red) stimuli (similar to Fig. 4h). Error bars depict the 95% confidence interval of the mean (2×s.e.m). Lines depict exponential fits, shading shows 95% prediction bounds of the fits. **f,** Top, projections of V1 activity during the visual stimulus presentation in control (black) and LM silencing (blue) trials onto the feedback communication direction calculated for the silencing window at time T5 (256 ms). V1 activity in different silencing windows (T4, T5 shown above, T6 and T7 below) is then projected onto the same feedback communication direction calculated for T5 during Go (left) and No-go trials (right). In No-go trials, the feedback direction at time T5 consistently separates control from silencing trials, not only at T5, but also at earlier and later silencing onset times. During Go trials this separation between control and silencing trials decreases rapidly with increasing time intervals. This indicates more rapid changes in feedback directions over time in Go, compared to No-go trials. Bottom, schematic illustrating the faster temporal re-orientation of the feedback communication direction in Go (green) as opposed to No-go (red) trials. **g,** Cross-validated cosine similarity matrices for the feedforward communication directions (influence of V1 silencing on LM population activity) calculated using an alternative method. Instead of LDA, the difference between the means of population activity in control and silencing trials (without normalizing to the covariance matrix, see Methods) was used to calculate communication directions. **h,** As in **g,** but for feedback communication directions (influence of LM silencing on V1 population activity). **i,** Initial decay slope of communication directions in Go (green) and No-go (red) trials, calculated from the difference between the means of population activity in control and silencing trials (corresponding to **g** and **h**) instead of using LDA. Error bars depict the 95% confidence interval of the mean (2×s.e.m). **j,** Relationship between the initial slopes (slopes between lag 0 and lag 1) of the decay over time lags of communication direction similarities and of activity direction similarities in the target area (left), and in the source area (right), calculated from the difference between the means of population activity in control and silencing trials (corresponding to **g** and **h**) instead of using LDA. Orange and purple dots show data from individual animals during V1 silencing (feedforward influences) and LM silencing (feedback influences) experiments, respectively. P-values from two-sided Wilcoxon signed-rank tests.

**Extended Data Fig. 6.**
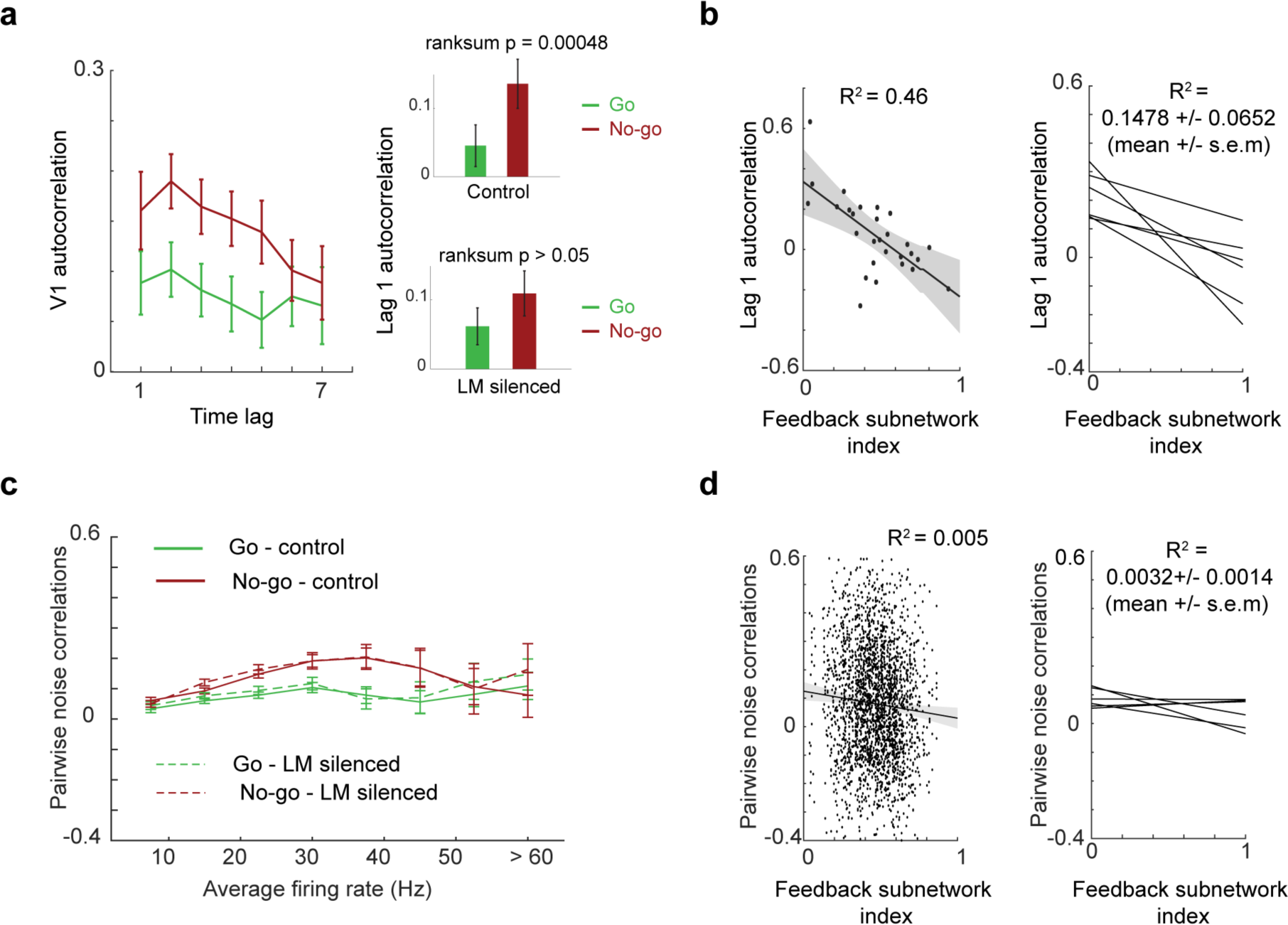
-Pairwise noise correlations and autocorrelations of V1 cells. **a,** Left, Autocorrelation of V1 cells as a function of time lags, calculated for each cell separately and averaged across cells during Go (green) and No-go (red) trials. Right, Autocorrelation of V1 neurons at time lag 1 (∼65ms time lag) during Go (green) and No-go (red) stimulus, in control trials (top) and during LM silencing (bottom). Error bars depict the 95% confidence interval of the mean (2×s.e.m). A comparison of autocorrelation strength in control and silencing trials is only possible at time lag 1, since silencing was not performed continuously but during short time windows. **b,** Left, Relationship between autocorrelation at time lag 1 (∼65ms time lag) and the feedback influence index of individual V1 neurons of one example mouse in Go trials. The feedback influence index quantifies how much a V1 neuron was influenced by feedback, calculated as its maximum coefficient in the feedback communication direction across the eight silencing time windows. Line represents the linear fit, shading depicts 95% prediction bounds of the fit. Right, linear fits as on the left for all mice. Each line represents a linear fit to the relationship between autocorrelation and feedback influence index of V1 neurons from one animal (n = 6). **c,** Average strength of noise correlations between pairs of V1 cells as a function of their average firing rates during Go (green) and No-go (red) stimuli, when LM was silenced (dashed lines) and in the corresponding time windows in control trials (solid lines). Error bars depict the 95% confidence interval of the mean (2×s.e.m). **d,** Left, relationship between pairwise noise correlation of pairs of V1 cells and the average feedback influence index of the pair of one example mouse in Go trials. Feedback influence index for each V1 neuron was calculated as in **b.** Right, linear fits as in **c** for all mice (n = 6).

